# Synthesis and antimicrobial evaluation of new cephalosporin derivatives containing cyclic disulfide moieties

**DOI:** 10.1101/2021.11.03.467114

**Authors:** Inga S. Shchelik, Karl Gademann

## Abstract

Due to a steady increase of microbial resistance, there is a need to increase the effectiveness of antibiotic performance by involving additional mechanisms of their penetration or retention for their better action. Cephalosporins are a successful group of antibiotics to combat pathogenic microorganisms, including drug-resistant strains. In this study, we investigated the effect of newly synthesized cephalosporin derivatives with cyclic disulfide modifications against several Gram-positive and Gram-negative strains as well as against biofilm formation. The incorporation of asparagusic acid was found to be effective in improving the activity of the drug against Gram-negative strains compared to the all carbon control compounds. Furthermore, we could demonstrate the successful reduction of biofilm formation for *Staphylococcus aureus* and *Pseudomonas aeruginosa* at similar concentrations as obtained against planktonic cells. We propose that the incorporation of cyclic disulfides is one additional strategy to improve antibiotic activity and to combat bacterial infections.

---

Over the last decades, the understanding of the relevant biological pathways and targets for antibiotic activity has been significantly expanded. Historically, activity on the bacterial target was most importantly optimized and has been constantly improved by molecular design,^1–3^ synthetic chemistry,^3–8^ and analytical techniques up to recently introduced cryo EM structure determination.^9^ Less studied but equally important, several additional factors strongly contribute to overall antibiotic performance, including bacterial uptake of antibiotics, their biotransformation to inactive substances, the active efflux of those agents by bacteria, and others.^10,11^ As a consequence, performance enhancement of antibiotics has recently enlarged to also include these aspects. Several strategies are currently employed to increase uptake of antibiotics. For example, the “Trojan horse” approach exploits receptor-mediated Fe-uptake mechanisms in bacteria, culminating in the launch of the cephalosporin-based antibiotic cefiderocol on the market.^12,13^ Alternatively, influx can be enhanced by the self-promoted uptake mechanism with polycationic antibiotics involving the displacement of the divalent cations that stabilize the lipopolysaccharides (LPS) membrane packing, thereby increasing the passage of the promoter into the periplasm.^14^ Moreover, some efforts were made to investigate the dithiol-mediated uptake of antibiotics by bacteria^15^ in a similar manner to eukaryotic cells,^16–19^ however the obtained results have been mixed so far. Another important aspect is the formation of biofilms, which renders the producing bacteria highly recalcitrant to antibiotics.^20^ Biofilms are produced upon switching of the bacterial growth mode from planktonic to sessile, where the bacteria aggregate and create a mesh-like structure with extracellular substance.^21^ In recent years, novel therapeutic strategies were developed, where different classes of antimicrobials are able to interfere at different levels with the formation of biofilms^22^, including disulfide containing compounds.^23,24^ In an earlier study reported by us, we demonstrated that introducing sulfur-containing moieties into the antibiotic vancomycin achieved potent antibacterial activity against resistant *Enterococcus faecalis, Enterococcus faecium,* and Gram-negative *Moraxella catarrhalis.* Moreover, the obtained derivatives featured improved activity against biofilm formation.^25^ In this study, we extended this concept to a series of cephalosporin derivatives with cyclic disulfide moieties and their full carbon analogs to closer investigate the potential of such modifications on biological activity of the antibiotic (Figure 1).

**Figure 1.**
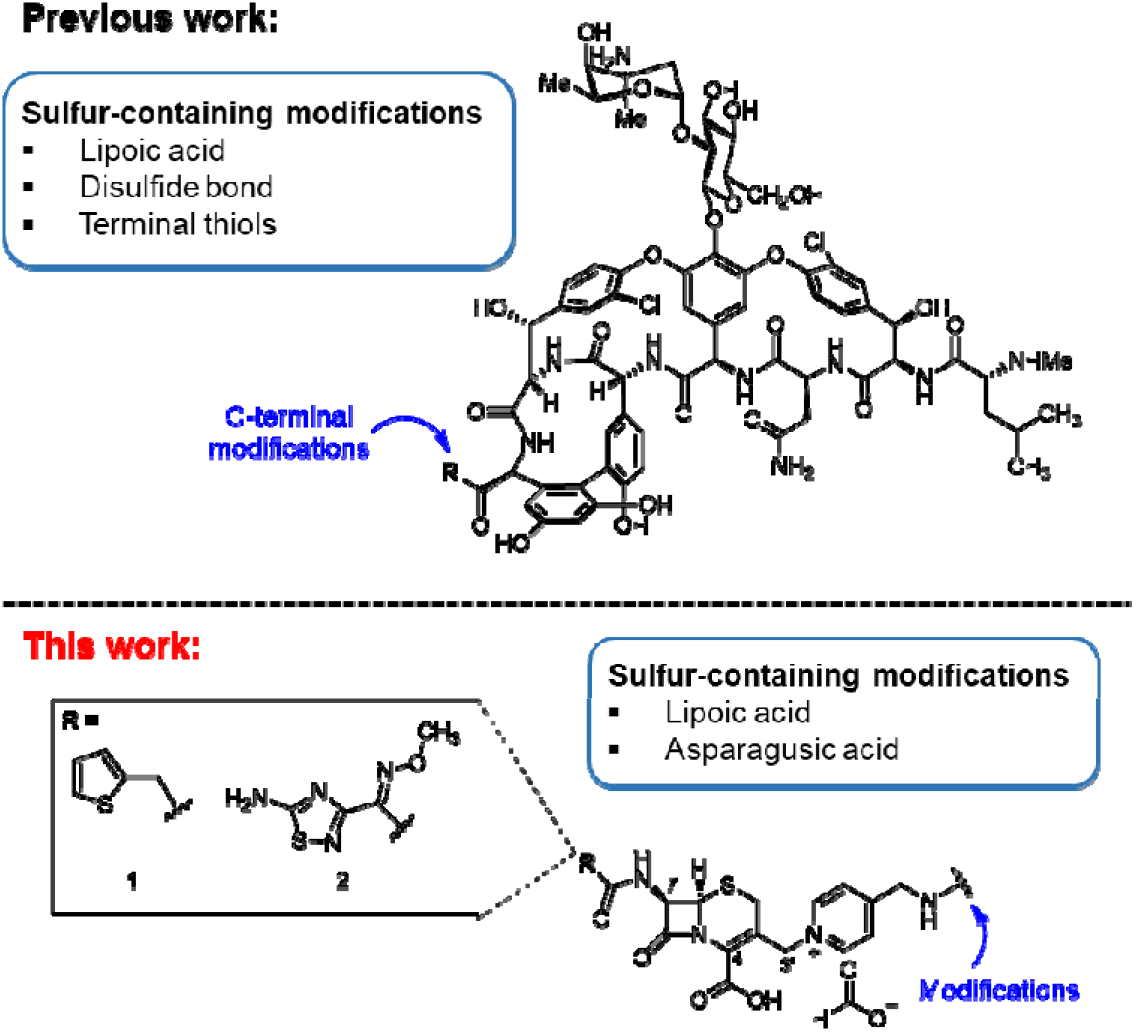
Sulfur-containing modification of different antibiotics. Top: Previous work on vancomycin modifications.^25^ Bottom: This work on cephalosporin modifications.

## Results

### Design

Cephalosporins are a class of *β*-lactam antibiotics originally derived from the mold *Emericellopsis minimum* (formerly *Cephalosporium acremonium*).^26,27^ Cephalosporins contain a six-membered dihydrothiazine ring attached to the *β*-lactam part and share the mechanism of action with *β*-lactam antibiotics. By mimicking the D-Ala-D-Ala site of peptidoglycan precursor, cephalosporins bind penicillin binding protein (PBP) thereby irreversibly inhibiting PBP crosslinking of peptidoglycan and disrupting bacterial cell wall structural integrity.^28^ The biological activity of these antibiotics can be varied depending on modifications at C3, C4, and C7 positions. In our research, thiophene (**1**) and amino-thiadiazol (**2**) based side chains were chosen for the variation of the C7-position of the cephalosporin core. The thiophene moiety is found in several examples of 1^st^ and 2^nd^ generation cephalosporins, which are mainly active against Gram-positive bacteria. On the other hand, the presence of amino-thiadiazol side chain represents a key structural unit for the 4^th^ and 5^th^ generation derivatives, which demonstrated an increased activity against Gram-negative strains.^29^ To introduce the zwitterionic properties in cephalosporin the modification at the C3’-position of the cepham ring was done with the incorporation of the quaternary ammonium moiety.^30^ Pyridinium substitution, found in several clinically used antibiotics of 1^st^ and 3^rd^ generation, was chosen using 4-(aminomethyl)pyridine, which is suitable for introducing the desired linkers via amide bond formation.

The synthesis of cephalosporin derivatives with the thiophene side chain started from cephalothin (**3**), which was treated with N-methyl-N-(trimethylsilyl) trifluoroacetamide (MSTFA) followed by the addition of trimethylsilyl iodide (TMSI). Dichloromethane was removed after 1 h of stirring and the pyridine derivatives **4**, **5**, **6**, **7**, or **8** dissolved in DMF were added to access the desired products. The derivative with pyridinium moiety **11** lacking the additional linkers attached was prepared as the control compound from its Boc-protected precursor **10** by the deprotection reaction in acidic conditions. The derivatives including lipoic and asparagusic acid moieties (**12b**, **13b**) were prepared for the closer investigation of possible dithiol-mediated uptake of cephalosporin by Gram-negative strains. Moreover, the full-carbon analogs (**12a**, **13a**) were also synthesized as control compounds to explore the necessity of disulfide bond incorporation for the improving of the obtained antibiotics activity. The reaction of cefalotin **3** with the linker **7** led to the formation of several by-products, which were not possible to separate from the desired compound. Therefore, the asparagusic acid derivative of cephalosporin with a thiophene side chain was obtained through intermediate **14**, which, after Boc deprotection and oxidation reactions, resulted in the formation of the desired product **13b** (Figure 2, A).

**Figure 2.**
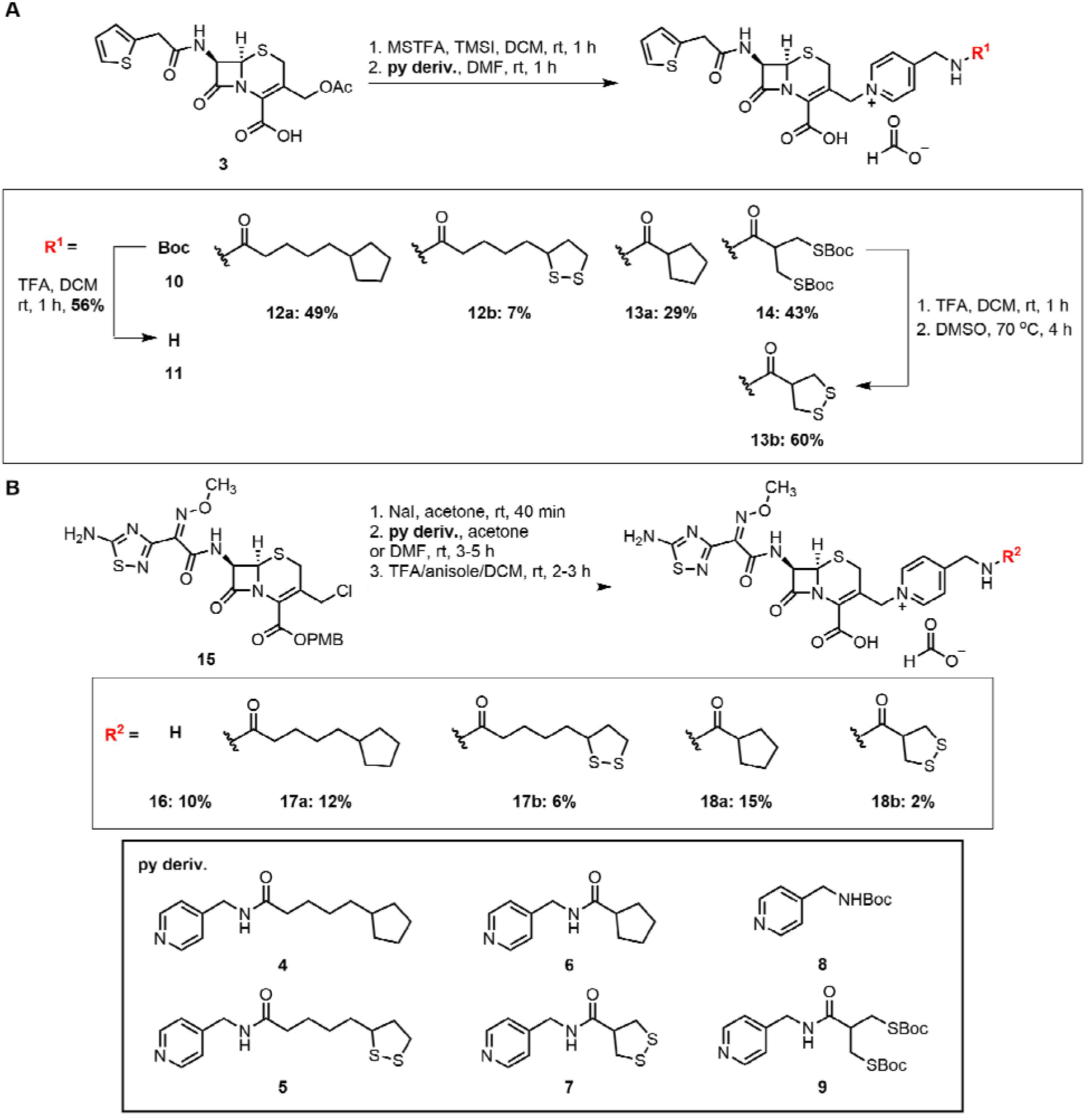
Functionalization of cephalosporins. A. Synthesis of cephalosporin derivatives containingthiophene side chain. B. Synthesis of cephalosporin derivatives containing amino thiadiazol side chain.

The synthesis of the desired derivatives of cephalosporin with an amino-thiadiazol side chain started from the substitution of cephalosporin core **15** with pyridine derivatives **4**, **5**, **6, 7** and **8** *via* three straightforward steps without the isolation of intermediates, leading to the scope of disulfide-containing products (**17b**, **18b**), their full-carbon analogs (**17a**, **18a**), and the control compound with only the pyridinium moiety attached (**16**) (Figure 2, B).

### Antibacterial activity

The antibacterial activity of obtained cephalosporin derivatives was determined against a selection of pathogenic and clinically isolated Gram-positive and Gram-negative strains by standard broth microdilution susceptibility tests.^31^ The data is summarized in Table 1 (more data are presented in Table S1-S2, Supporting Information).

**Table 1.**
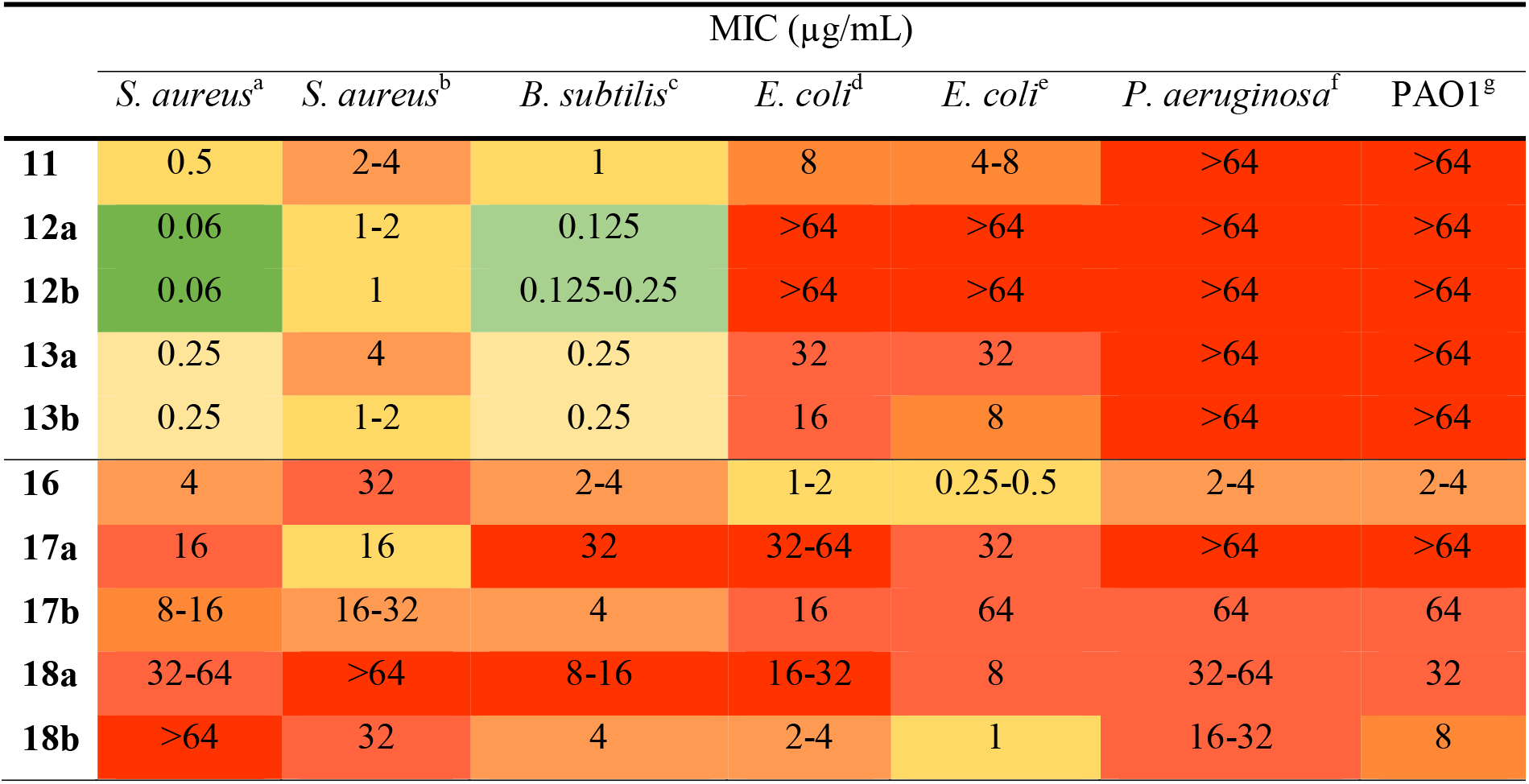
Antimicrobial activity of cephalosporin derivatives. Compounds **XXa** are featuring the full-carbon control compounds and compounds **XXb** are cyclic disulfides. ^a^ATCC 25923, ^b^ATCC 43300, ^c^ATCC 6633, ^d^ATCC 25922, ^e^K12 MG1622, ^f^ATCC 27853, *^g^P. aeruginosa* PAO1.

Thiophene-containing cephalosporin derivatives featuring elongated side chains (**12** and **13**) displayed an increase in antibacterial activity when tested against *Staphylococcus aureus (S. aureus)* and *Bacillus subtilis (B. subtilis)* sensitive strains compared to control compound **11**. In contrast, loss in activity was observed for the derivative with lipoic acid **12a** and its full-carbon analog **12b** against Gram-negative *Escherichia coli (E. colĩ).* No activity was detected for the obtained derivatives against *Pseudomonas aeruginosa (P. aeruginosa),* which fully correlated with the data for previously reported cephalosporin derivatives with thiophene modification at the C-7-position of the cepham core.^32^ The activity of cephalosporin derivatives of the amino-thiadiazol side-chain series was mostly detected for Gram-negative strains (see control compound **16**). However, the incorporation of lipoic acid-based side chain significantly decreased the activities against *E. coli* and *P. aeruginosa.* The similar effect was observed for its full-carbon analog **17a**. On the other hand, an 8-fold higher activity was found for the asparagusic acid-containing compound **18b** compared to its non-disulfide analog **18a** when tested against *E. coli* and almost no loss in activity compared to the control compound **16** (Table 1). A strong decrease in activity was observed among the compounds with substitutions on pyridinium linker (**17a**, **17b**, **18a**) against *Pseudomonas* strains, compared to the control compound **16**. Exceptionally, only a 2-4-fold decline in activity was observed for the compound **18b** with the asparagusic acid modification against *P. aeruginosa* PAO1 when compared to parent compound **16**.

### Antibiofilm formation activity

Approximately 80% of pathogens that form biofilms are associated with persistent infections. The typical examples of such pathogens are *P. aeruginosa*, which is associated with cystic fibrosis,^33^ and *S. aureus,* which is responsible for wound infections.^34^ There are several examples of natural and synthetically obtained disulfide-containing antibacterials displaying the activity against biofilm formation. For example, the studies of methyl disulfide^24^ showed the inhibitory effect of the compounds against *Mycobacterium smegmatis* biofilm formation. Moreover, diphenyl disulfides^35^ or diallyl disulfides^36^ exhibited significant reduction of biofilm formation in *Pseudomonas* strains related to quorum sensing inhibition.^37^ Therefore, in our work, we decided to investigate the influence of the disulfide bond present in the structure of newly obtained cephalosporin derivatives on the biofilm formation of several Gram-positive and Gram-negative strains.

#### Assessment of biofilm biomass

Earlier in our group, the biofilm-forming activity of six Gram-positive strains was investigated in different media. Strong biofilm formation was observed for *S. aureus* and *E. faecalis* strains in Brain Heart Infusion media with the addition of 1% glucose.^25^ In this study, we focused on the biofilm formation of Gram-negative bacteria. *E. coli* and *Pseudomonas* were grown in eight different media: MHB, TS, TSG, TS2G, BHI, BHIG, M63, M63A for 24 h.^38^ The assessment of biofilm production was carried out by a crystal violet (CV) staining.

From the obtained results, biomass as measured by the absorbance of CV at 570 nm displayed low values for *E. coli* strains (lower 0.5) in all tested mediums. *Pseudomonas* featured excellent biofilm formation in M63 medium reported earlier.^39^ The replacement of glucose and casamino acids in M63 medium by arginine (M63A) initiated the better biofilm formation. However, the absorbance of CV measured in M63A medium for PAO1 was relatively low in comparison to *P. aeruginosa* ATCC 27853, with values ranging from 1.0 to 1.6, in contrast to 5.1 to 6.9, respectively (Figure 3). From the obtained results *P. aeruginosa* ATCC 27853 strain was chosen as a strong biofilm producer in M63 medium supplemented with arginine for the further investigation of antibiofilm formation activity of newly obtained cephalosporin derivatives.

**Figure 3.**
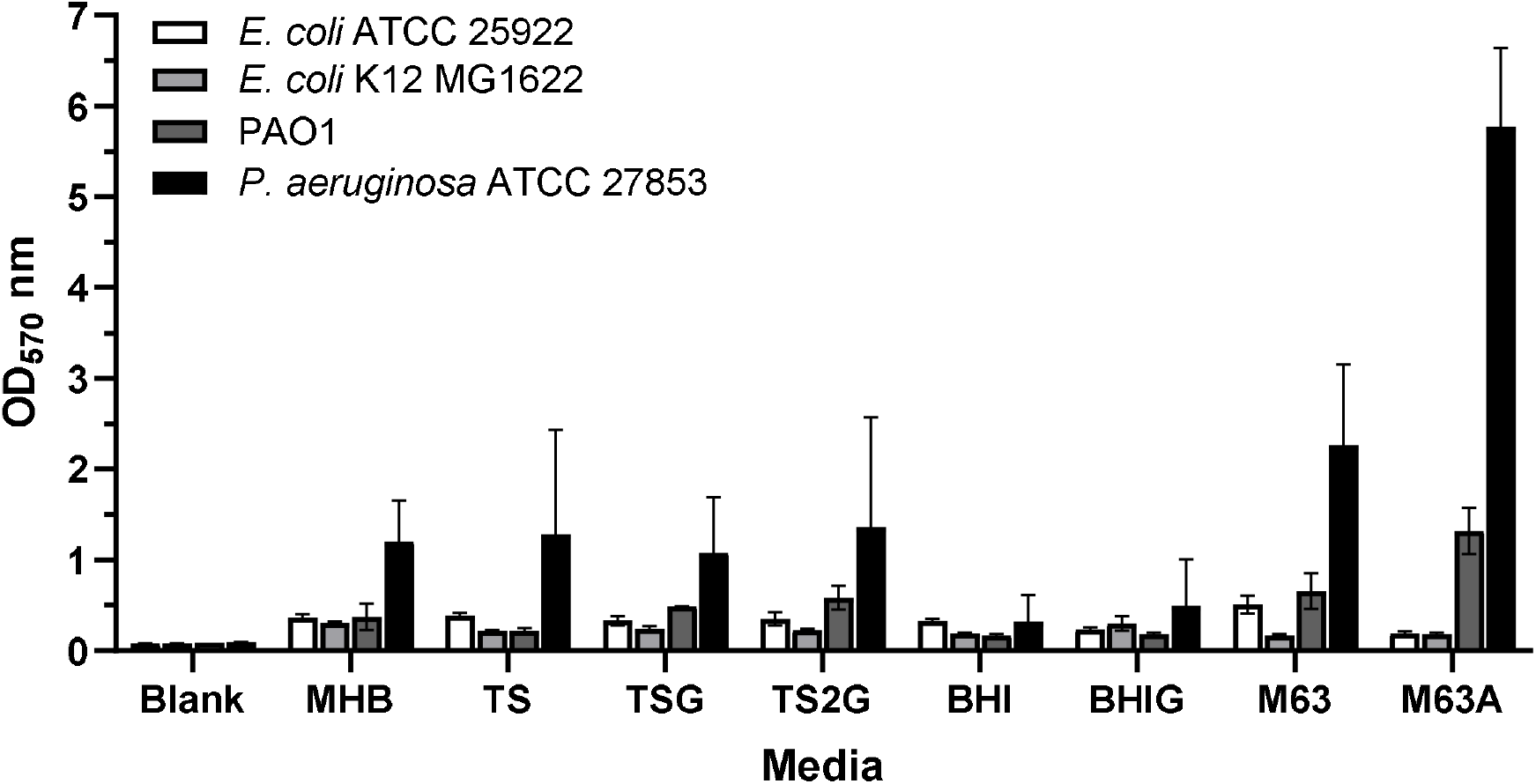
Assessment of biofilm biomass by crystal violet staining of four Gram-negative strains: *E. coli* ATCC 25922, *E. coli* K12 MG1622, PAO1, *P. aeruginosa* ATCC 27853. Strains were incubated in eight different media. Error bars represent standard deviation of two independent experiments in triplicate.

#### Assessment of metabolic activity of biofilm cells by fluorescein diacetate (FDA)

For the assessment of metabolic activity of the *Pseudomonas* biofilms the fluorescein diacetate (FDA) was chosen as an excellent dye when tested against Gram-negative strain.^40^ Biofilms were formed in 24 h before FDA assay. The formed biofilms were treated with FDA solutions at concentrations 2, 4 and 8 μg/mL and the plates were incubated at 37 °C. The relative fluorescence units (RFU) were taken every 2 min for 2 h. *P. aeruginosa* showed increase in RFU values during 120 min (Figure S1 in the Supporting Information). The selection of optimal assay conditions was based on calculated statistical quality parameters (Table S3 in the Supporting Information). The excellent Z’ value was obtained only at concentration 2 μg/mL, the high variability was observed between replicates at concentrations 4 or 8 μg/mL. According to the obtained results the conditions for the FDA assay are 2 μg/mL with incubation for 60 min for *P. aeruginosa* ATCC 27853.

#### MBRC determination

Using the previously optimized assay, novel synthesized cephalosporin derivatives were tested for their ability to repress the growth and biofilm formation of four bacterial strains. The biofilms were grown in optimized conditions in the presence of antibiotics at different concentrations. After incubation, the metabolic activity of formed biofilms was determined by resazurin for Gram-positive strains^25^ or FDA for *P. aeruginosa* with the chosen conditions. Colony forming unit (CFU) was also defined to validate the results obtained with metabolic activity assessment assays.

Within cephalosporin derivatives, 1^st^ generation analogs showed no significant difference in MBRC values compared to MIC results when tested against *Staphylococcus* strains. Moreover, compounds containing disulfide bond exhibited similar results compared to the control full-carbon derivatives. The same trend was observed for the 4^th^ generation cephalosporin derivatives when tested against Gram-positive strains (Table S4, Supporting Information). However, 4-fold enhancement in activity was detected for the compound with asparagusic acid moiety **18b** compared to its non-disulfide analog **18a** when tested against *P. aeruginosa* biofilm formation (Table 2). These results demonstrate the positive influence of the disulfide bond on biological activity of the modified antibiotic when tested against the biofilm formation of Gram-negative bacteria.

**Table 2.**
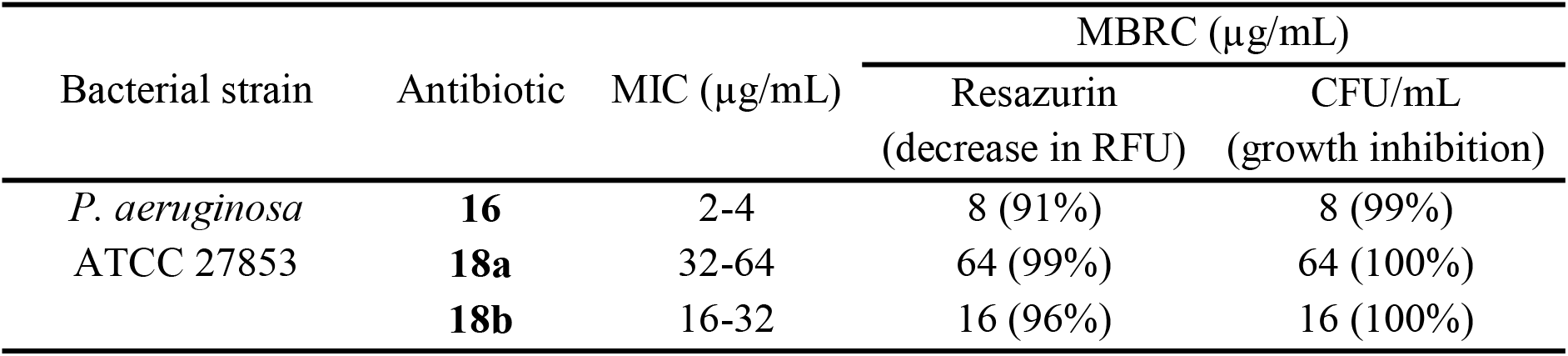
Minimal biofilm reduction concentration (MBRC) determined by resazurin and cell counting methods

## Discussion

Performance enhancement of antibiotics *via* facilitating drug uptake by bacteria remains an important challenge. In this study, we could demonstrate the increase of antibacterial activity against Gram-negative strains for the new cephalosporin derivative containing asparagusic acid moiety and an amino thiadiazol side chain (**18b**). We hypothesized that the introduction of disulfide bond led to either better penetration of antibiotic through the outer membrane or involving additional mechanism of action. The benefit of disulfide bond presence in the structure of an antibiotic was earlier demonstrated by Nicolaou and co-workers, where they showed that the removal of disulfide bond from the natural product psammaplin A or its analogs dramatically reduces the activity of the compound against a range of pathogenic strains.^41^

An earlier study by the group of Matile reported loss of activity of several antibiotics against Gram-negative strains after the introduction of cyclic oligochalcogenides into their structure.^15^ In this study, a similar effect was observed after the incorporation of lipoic acid into the cephalosporin structure (**12b**, **17b**). However, a similar tendency was observed for the control compounds with full-carbon moieties (**12a**, **17a**). These results suggest that the incorporation of an additional aliphatic chain as a structural modification of the antibiotic was not optimal. The increased size of the compounds resulted in the loss of activity against Gram-negative strains, regardless of the presence of the disulfide bond. In contrast, an excellent activity against Grampositive strains was shown for the cephalosporin derivatives with a thiophene side chain after the modifications while remaining effective in reducing of biofilm layer formation as well. Additionally, against Gram-negative strains, an improvement in activity was observed for the cephalosporin derivative with amino thiadiazol side chain and the asparagusic acid modification (**18b**) when tested against *Pseudomonas.* However, as control compound **16** still displayed better performance when tested against Gram-negative strains, further optimization of the disulfide-containing antibiotics will be required. For example, the incorporation of different cyclic oligochalcogenides might potentially increase the antibiotic uptake and lower the amount of the drug needed for the reduction of biofilm formation and inhibition of bacterial growth. The mechanism of thiol-mediated uptake remains under scientific debate for eukaryotic systems,^16^ and for prokaryotes, even less is known at the moment.

### Conclusion

In summary, these results demonstrate that cyclic disulfides can lead to improved activity compared to their all carbon-control compounds. Moreover, cyclic disulfides apparently are able to overcome the outer membrane of bacteria or these antibiotics might invoke an additional mechanism of action. Together with data obtained in previous research done in our group on vancomycin,^25^ these results clearly demonstrate the usefulness of disulfides in overcoming resistance in bacteria and biofilms compared to the all carbon control compounds, and therefore, potentially improving antibiotic effectiveness against certain difficult to treat pathogens.

## Methods

### Chemistry

The reactions were carried out under inert gas (N_2_ or Ar) in oven-dried (120 °C) glass equipment and monitored for completion by TLC or UHPLC-MS (ESI). The chemicals and solvents for the reactions and analyses were obtained from the commercial sources and used without additional purification. DL-α-lipoic acid, 7-ACA, GCLE, and thiophene-2-acetyl chloride, ATDA were purchased from Sigma-Aldrich, Apolo Scientific, TCI, and ABCR, respectively. The synthesis of cephalothin **3**, cephalosporin derivative **15**, cephalosporin derivative **16**, 5-cyclopentylpentanoic acid, and 3-(acetylthio)-2-((acetylthio)methyl)propanoic acid were carried out following literature procedures.^42–46^ The synthesized compounds were characterized through ^1^H-NMR and ^13^C NMR spectroscopy using Bruker Avance 500 or 400 spectrometers and CDCl_3_, MeOD, D_2_O, or DMSO-*d*_6_ as the solvents. High-resolution mass spectra (HRMS) were obtained by using timsTOF Pro TIMS-QTOF-MS instrument in the ESI positive mode, and the values were expressed in *m/z.* The synthesized compounds were purified either through column chromatography using glass columns packed with silica gel 60 (230-400 Mesh) purchased from Sigma-Aldrich with the solvent mixture indicated or reverse phase HPLC. Thin Layer Chromatography (TLC) was run on Merck TLC plates silica gel 60 F254 on glass plates with the indicated solvent system; the spots were visualized by UV light (365 nm), and stained by anisaldehyde, ninhydrin, or KMnO_4_ stain. Additional information about the equipment used can be found in the Supporting Information.

### Procedures for the synthesis of linkers for cephalosporin derivatization

#### Synthesis of N-(pyridin-4-ylmethyl)cyclopentanecarboxamide

Cyclohexane carboxylic acid (0.191 mL, 1.76 mmol) was dissolved in dry DCM (3 mL) and the solution was cooled down to 0 °C. A solution of oxalylchloride (0.272 mL, 3.17 mmol) in dry DCM (3 mL) was added dropwise. The reaction mixture was allowed to warm to room temperature and was stirred for 3 hours. The solution was concentrated at reduced pressure to afford a slightly yellow solid, which was used for the next step without further purification.

The obtained cyclopentanecarbonyl chloride (115mg, 0.867 mmol) was dissolved in dry DMF (4 mL), 4-(aminomethyl)pyridine (0.100 mL, 0.960 mmol) was added dropwise followed by the addition of DIPEA (distilled, 0.190 mL, 1.15 mmol). The reaction mixture was stirred for 2 h and then poured into water. The aqueous phase was extracted with EtOAc (3×20 mL), the organic phase was washed with brine, dried over Na_2_SO_4_, and concentrated under reduced pressure. Purification was carried out by column chromatography (DCM:MeOH 95:5). The product was obtained as an amorphous solid (140 mg, 0.867 mmol, 79%). **R*f*** = 0.32 (DCM:MeOH 95:5). **^1^H NMR** (500 MHz, CDCl_3_) δ 8.53 (m, 2H), 7.17 (m, 2H), 6.03 (s, 1H), 4.45 (d, *J* = 5.5 Hz, 2H), 2.59 (p, *J* = 7.9 Hz, 1H), 1.98 – 1.67 (m, 6H), 1.59 (m, 2H). **^13^C NMR** (126 MHz, CDCl3) δ 176.62, 150.08, 147.96, 122.37, 45.85, 42.40, 30.63, 26.07. **IR (film):** 3287, 2955, 1647, 1603, 1539, 1416, 1239, 794. **HRMS (ESI):** calcd for C12H17ON2 [M+H]^+^, *m/z* = 205.13354, found 205.13343.

#### Synthesis of 3-(tritylthio)-2-[(tritylthio)methyl]propanoic acid

Triphenylmethane thiol (1.410 g, 5.1 mmol) was suspended in DMSO (3 mL) and DBU (1 mL, 6.12 mmol) was added. The reaction mixture turned dark orange and then a precipitate was observed. After the stirring of the reaction mixture at rt for 5 min, 3-bromo-2-(bromethyl)propionic acid (500 mg, 2.04 mmol) in 1 ml of DMSO was added dropwise. The precipitate dissolved over 2.5 h while stirring at rt. A 5% solution of HCl in water (20 mL) was added dropwise to the reaction and the water phase was extracted with DCM (3×15 mL). The organic phase was collected, dried over Na_2_SO_4_, and concentrated by reduced pressure. The crude mixture was purified by column chromatography (pentane:EtOAc:0.1%TFA 4:1, 2:1, 1:1). The desired product was isolated as a white solid (715 mg, 2.04 mmol, 55%).

**R*f*** = 0.8 (pentane:EtOAc = 1:1). **^1^H NMR** (500 MHz, CDCl_3_) δ 7.35 – 7.30 (m, 12H), 7.24 – 7.21 (m, 12H), 7.20 – 7.14 (m, 6H), 2.39 (dd, *J* = 12.5, 7.5 Hz, 2H), 2.19 (dd, *J* = 12.5, 5.9 Hz, 2H), 1.90 (p, *J* = 6.4 Hz, 1H). ^13^**C NMR** (126 MHz, CDCl3) δ 144.56, 129.71, 128.05, 126.85, 67.15, 32.79. **IR (film):** 1705, 1488, 1184, 1034, 908, 741, 699 **HRMS (ESI):** calcd for C_42_H_35_O_2_S_2_ [M+H]^+^, *m/z* = 635.20840, found 635.20873.

#### Synthesis of N-(pyridin-4-ylmethyl)-3-(tritylthio)-2-((tritylthio)methyl)propanamide

To a solution of 3-(tritylthio)-2-[(tritylthio)methyl]propanoic acid (174 mg, 0.273 mmol) in DMF (2.6 mL), PyBoP (284 mg, 0.546 mmol), HOBt (37 mg, 0.273 mmol), and DIPEA (0.1 mL, 0.55 mmol) were added. The reaction mixture was stirred for 2 h at rt. Water (20 mL) was added to the mixture and the aqueous layer was extracted with DCM (3×30 mL). The collected organic phases were dried over Na_2_SO_4_ and concentrated under reduced pressure. Purification was carried out by column chromatography (pentane:EtOAc 1:1, 1:2). The product was isolated as a white amorphous solid (184 mg, 0.273 mmol, 93%).

**R*f*** = 0.4 (pentane:EtOAc = 1:2). **^1^H NMR** (500 MHz, CDCl3) δ 8.45 (dd, *J* = 1.5, 4.5 Hz, 2H), 7.38 – 7.33 (m, 12H), 7.30 – 7.25 (m, 12H), 7.24 – 7.19 (m, 8H), 5.42 (t, *J* = 6.1 Hz, 1H), 4.38 (d, *J* = 6.1 Hz, 2H), 2.47 (dd, *J* = 12.5, 8.6 Hz, 2H), 2.17 (dd, *J* = 12.5, 5.6 Hz, 2H), 1.36 (tt, *J* = 8.6, 5.6 Hz, 1H). **^13^C NMR** (126 MHz, CDCl3) δ 172.58, 149.98, 144.62, 129.71, 128.09, 126.94, 122.49, 67.20, 66.24, 46.77, 42.40, 33.88. **IR (film):** 1663, 1600, 1539, 1488, 1443, 1415, 1387, 1318, 1265, 1183, 1096, 1033, 1000, 841. **HRMS (ESI):** calcd for C_48_H_43_ON_2_S_2_ [M+H]^+^, *m/z* = 727.28113, found 727.28012.

#### Synthesis of O,O’-di-tert-butyl S,S’-(2-((pyridin-4-ylmethyl)carbamoyl)propane-1,3-diyl) bis(carbonothioate)

N-(pyridin-4-ylmethyl)-3-(tritylthio)-2-((tritylthio)methyl)propanamide (170 mg, 0.234 mmol) was dissolved in DCM (1.2 mL) and TFA (1.2 mL, v/v 50%) was added. The reaction mixture turned yellow. TES (0.380 mL, 2.34 mmol) was added and the reaction mixture was stirred at rt under argon for 1 h. The solvent was evaporated and the mixture was washed with pentane (5×10 mL) until the white precipitate was completely dissolved. The deprotected product as a colorless oil was used for the next step without additional purification.

The dithiol derivative was next dissolved in CH_3_CN (2.4 mL). K_2_CO_3_ (65 mg, 0.470 mmol) was added, followed by the addition of a solution of Boc_2_O (0.100 mL, 0.490 mmol) in CH_3_CN (0.2 mL). The reaction was stirred at rt under argon for 5 h. The reaction mixture was concentrated under reduced pressure and DCM and water were added. The aqueous layer was extracted with DCM (3×15 mL). The organic phase was washed with brine and dried over Na_2_SO_4_. The solvent was evaporated and the residue was purified by column chromatography (pentane:EtOAc 1:1, 1:2). The product was isolated as a yellowish amorphous solid (66 mg, 0.234 mmol, 64%).

**R*f*** = 0.25 (pentane:EtOAc = 1:2). **^1^H NMR** (400 MHz, MeOD) δ 8.50 – 8.44 (m, 2H), 7.43 – 7.38 (m, 2H), 4.45 (s, 2H), 3.35 (s, 1H), 3.10 (dd, *J* = 13.8, 5.6 Hz, 2H), 3.00 (dd, *J* = 13.5, 8.6 Hz, 2H), 2.93 – 2.85 (m, 1H), 1.49 (s, 18H). **^13^C NMR** (101 MHz, MeOD) δ 174.88, 169.83, 150.61, 149.99, 123.96, 86.24, 43.06, 33.36, 28.44. **IR (film):** 2980, 2932, 1716, 1700, 1646, 1604, 1455, 1416, 1394, 1369, 1200, 1119, 833. **HRMS (ESI):** calcd for C_20_H_31_O_5_N_2_S_2_ [M+H]^+^, *m/z* = 443.16689, found 443.16715.

#### Synthesis of S,S’-(2-((pyridin-4-ylmethyl)carbamoyl)propane-1,3-diyl) diethanethioate

Oxalyl chloride (94 ul, 1.1 mmol) was added dropwise at 0 °C into the solution of 3-(acetylthio)-2-((acetylthio)methyl)propanoic acid (274 mg, 1.16 mmol) in DCM (11.5 mL) in presence of catalytic amount of N,N-dimethylformamide. The reaction mixture was stirred at 0 °C for 2 h. The solvent was evaporated under reduced pressure and the product was used for the next step without additional purification.

The obtained acid chloride (148 mg, 0.581 mmol) was dissolved in dry DMF (5.8 mL). 4-(aminomethyl)pyridine (0.065 mL, 0.6 mmol) was added dropwise into the reaction at rt, followed by the addition of DIPEA (distilled, 108 ul, 0.639 mmol). The reaction mixture was stirred at rt for 1.5 h. The solvent was evaporated and the residue was purified by column chromatography (DCM:MeOH = 100:5). The desired product was isolated as a yellow solid (80 mg, 0.581 mmol). **R*f*** = 0.35 (DCM:MeOH = 100:5). **^1^H NMR** (500 MHz, CDCl3) δ 8.58 – 8.54 (m, 2H), 7.25 – 7.21 (m, 2H), 6.46 (t, *J* = 6.1 Hz, 1H), 4.48 (d, *J* = 6.0 Hz, 2H), 3.16 (dd, *J* = 13.8, 7.4 Hz, 2H), 3.08 (dd, *J* = 13.8, 6.2 Hz, 2H), 2.67 – 2.58 (m, 1H), 2.34 (d, *J* = 1.0 Hz, 6H). **^13^C NMR** (126 MHz, CDCl3) δ 196.14, 172.19, 150.03, 147.38, 122.50, 47.39, 42.62, 30.77. **IR (film):** 3301, 1690, 1603, 1542, 1416, 1355, 1249, 1133, 957, 788, 625. **HRMS (ESI):** calcd for C14H19N2O3S2 [M+H]^+^, *m/z* = 327.08316, found 327.08286.

#### Synthesis of *N-(pyridin-4-ylmethyl)-1,2-dithiolane-4-carboxamide*

S,S’-(2-((pyridin-4-ylmethyl)carbamoyl)propane-1,3-diyl) diethanethioate (100 mg, 0.306 mmol) was dissolved in MeOH (3.06 mL) and K_2_CO_3_ (127 mg, 1 mmol) was added. The reaction mixture was stirred at rt for 1 h. The solvent was evaporated and water (20 mL) was added. The water phase was extracted with EtOAc (3×20 mL). The organic phase was collected, washed with brine, dried over Na_2_SO_4_, and concentrated by reduced pressure. The crude was dried under vacuo and then dissolved in DMSO (1 mL). The reaction mixture was heated to 70 °C and stirred for 12 h in open flask. The solvent was evaporated and the reaction mixture was purified by column chromatography (DCM:MeOH= 90:10). The desired product was obtained as a slightly yellow amorphous solid (21 mg, 0.306 mmol, 29%).

**R*f*** = 0.6 (DCM:MeOH = 90:10). **^1^H NMR** (500 MHz, CDCl3) δ 8.60 – 8.51 (m, 2H), 7.21 – 7.13 (m, 2H), 6.42 (s, 1H), 4.47 (d, *J* = 6.0 Hz, 2H), 3.45 – 3.38 (m, 4H), 3.37 – 3.32 (m, 1H). **^13^C NMR** (126 MHz, CDCl3) δ 172.17, 150.31, 147.03, 122.35, 52.39, 42.84. **IR (film):** 2925, 2370, 1644, 1602, 1562, 1445, 1415, 1363, 1315, 1260, 1219, 1030, 994, 796, 609, 476. **HRMS (ESI):** calcd for C_10_H_13_ON_2_S_2_ [M+H]^+^, *m/z* = 241.04638, found 241.04626.

#### Synthesis of 5-cyclopentyl-N-(pyridin-4-ylmethyl)pentanamide

To a solution of 5-cyclopentylpentanoic acid (40 mg, 0.235 mmol) in DMF (2.40 mL), HATU (179 mg, 0.47 mmol) and DIPEA (distilled, 0.120 mL, 0.705 mmol) were added. The reaction mixture turned slightly brown. After 5 min of stirring, 4-(aminomethyl)pyridine (0.028 mL, 0.282 mmol) was added. The reaction mixture was stirred at room temperature for 1.5 h. Water (10 mL) was added and the mixture was extracted with EtOAc (4×20 mL). The combined organic layer was washed with brine, dried over MgSO4 and concentrated under reduced pressure. The purification carried out by column chromatography (pentane:EtOAc 1:1). The desired product was isolated as a white solid (46 mg, 0.235 mmol, 75%).

**R*f*** = 0.35 (pentane:EtOAc = 1:1). **^1^H NMR** (500 MHz, CDCl3) δ 8.55 (d, *J* = 5.8 Hz, 2H), 7.18 (d, *J* = 6.0 Hz, 2H), 6.02 – 5.80 (br, 1H), 4.46 (d, *J* = 6.1 Hz, 2H), 2.26 (t, *J* = 7.5 Hz, 2H), 1.80 – 1.62 (m, 4H), 1.62 – 1.54 (m, 2H), 1.53 – 1.46 (m, 2H), 1.40 – 1.27 (m, 5H), 1.13 – 0.99 (m, 2H). **^13^C NMR** (126 MHz, CDCl3) δ 173.48, 150.11, 147.79, 122.45, 42.43, 40.11, 36.85, 35.98, 32.82, 28.60, 26.10, 25.30. **IR (film):** 3284, 2926, 2857, 1652, 1528, 1351, 842, 558, 464. **HRMS (APCI):** C_16_H_25_ON_2_ [M+H]^+^, *m/z* = 261.19614, found 261.19591.

#### Synthesis of 5-(1,2-dithiolan-3-yl)-N-(pyridin-4-ylmethyl)pentanamide

To a solution of racemic α-lipoic acid (400 mg, 1.94 mmol) in dry DMF (18 mL), PyBOP (1.8 g, 3.52 mmol), HOBt (119 mg, 0.88 mmol), and DIPEA (1 ml, 5.28 mmol) were added. The reaction mixture was stirred for 10 min at rt followed by an addition of 4-(aminomethyl)pyridine (0.178 ml, 1.76 mmol). The reaction mixture was stirred overnight at rt and then the solvent was removed under reduced pressure. The purification was carried out by column chromatography (DCM:MeOH = 95:5). The desired product was isolated as a yellow oil (435 mg, 1.47 mmol, 83%). ***Rf*** = 0.35 (DCM: MeOH = 95:5) **^1^H NMR** (500 MHz, CDCl3) δ 8.55 (dd, *J* = 1.5, 4.5 Hz, 2H), 7.20 (dd, *J* = 1.5, 4.5 Hz, 2H), 5.91 – 5.85 (m, 1H), 4.47 (d, *J* = 6.1 Hz, 2H), 3.61 – 3.54 (m, 1H), 3.21 – 3.08 (m, 2H), 2.50 – 2.43 (m, 1H), 2.31 – 2.25 (m, 2H), 1.94 – 1.86 (m, 1H), 1.80 – 1.61 (m, 4H), 1.56 – 1.42 (m, 2H). **^13^C NMR** (126 MHz, CDCl3) δ 173.02, 150.05, 147.76, 122.50, 56.54, 42.50, 40.41, 38.64, 36.45, 34.73, 29.04, 25.45. **IR (film)**: 3284, 3054, 2926, 2855, 1650, 1602, 1543, 1416, 1361, 1259, 1067, 1028, 792 **HRMS (ESI)**: calcd for C_14_H_21_ON_2_S_2_ [M+H]^+^, *m/z* = 297.10898, found 297.10870.

### Procedures for the synthesis of cephalosporin derivatives with thiophene side chain

#### General procedure

Under an argon atmosphere, cephalothin (**3**, 1 equiv.) and MSTFA (3 equiv.) were added to dry methylene chloride (0.1 M solution) in a Schlenk tube, and the reaction mixture was stirred at rt for 15 min until a solution was observed. Iodotrimethylsilane (2 equiv.) was added and stirring was continued for another 1 h. The mixture was concentrated under reduced pressure connected to Schlenk line, and the residual oil was dissolved in dry DMF to obtain a 0.1 M solution. A few drops of dry THF were added and the mixture was stirred for 1 minute. The pyridine derivative dissolved in dry DMF (0.1 M solution) was added and the reaction mixture was stirred at rt for 1 h. The solvents were evaporated, and the residue was resuspended in a mixture of MeOH/CH_3_CN, filtered through a SPE column, and purified by preparative RP-HPLC to afford the desired products. The purification methods, analytical data, and yields are reported below.

#### Cephalosporin derivative **10**

RP-HPLC (Gemini-NX, flow rate = 15 mL/min): Gradient 5% B for 14 min; 0% - 50% B for 46 min; 50% - 100% B for 3 min, wash. The desired product, eluting at 28.5 min, was collected and lyophilized to afford product **10** (21 mg, 0.126 mmol, 31%) as a white solid. **_1_H NMR** (500 MHz, DMSO-*d*_6_) δ 9.40 (d, *J* = 6.5 Hz, 2H), 9.05 (d, *J* = 8.4 Hz, 1H), 7.94 (d, *J* = 6.4 Hz, 2H), 7.72 (t, *J* = 5.8 Hz, 1H), 7.35 – 7.33 (dd, *J* = 5.1, 1.3 Hz, 1H), 6.92 (dd, *J* = 5.1, 3.4 Hz, 1H), 6.89 (dq, *J* = 3.4, 1.1 Hz, 1H), 5.63 (d, *J* = 13.4 Hz, 1H), 5.54 (dd, *J* = 8.5, 4.8 Hz, 1H), 5.04 – 4.99 (m, 2H), 4.41 (d, *J* = 5.9 Hz, 2H), 3.74 – 3.67 (m, 2H), 3.51 (d, *J* = 17.6 Hz, 1H), 3.00 (d, *J* = 17.5 Hz, 1H), 1.41 (s, 9H). **^13^C NMR**(126 MHz, DMSO-*d*_6_) δ 169.89, 162.35, 155.86, 144.70, 136.96, 126.57, 126.22, 125.16, 124.93, 78.75, 60.99, 58.95, 57.34, 42.91, 40.43, 35.76, 28.14, 24.29. **HRMS (ESI)**: calcd for C25H29O6N4S2 [M^+^], *m/z* = 545.15230, found 545.15265. The purity of the compound was analyzed by analytical RP-HPLC (Gemini-NX). Gradient starts from 5% B for 4 min, 5% - 100% B for 11 min, wash. The product eluted at 11.6 min was detected at λ= 270 nm.

#### Cephalosporin derivative **12a**

RP-HPLC (Gemini-NX): Gradient 5% B for 14 min; 5% - 40% B for 46 min; 40% - 100% B for 5 min, wash. The desired product eluting at 48.2 min was collected and lyophilized to afford the product **12a** (17 mg, 0.057 mmol, 49%) as a white solid. **^1^H NMR** (500 MHz, MeOD) δ 9.10 (d, *J* = 6.6 Hz, 2H), 7.93 (d, *J* = 6.6 Hz, 2H), 7.25 (dd, *J* = 5.1, 1.1 Hz, 1H), 6.97 – 6.90 (m, 2H), 5.69 (d, *J* = 5.0 Hz, 1H), 5.67 (d, *J* = 14.1 Hz, 1H), 5.18 (d, *J* = 14.1 Hz, 1H), 5.07 (d, *J* = 5.0 Hz, 1H), 3.83 – 3.74 (m, 2H), 3.58 (d, *J* = 17.8 Hz, 1H), 3.09 (d, *J* = 17.8 Hz, 1H), 2.34 (t, *J* = 7.6 Hz, 2H), 1.74 – 1.82 (m, 3H), 1.69 – 1.58 (m, 4H), 1.58 – 1.50 (m, 2H), 1.42 – 1.32 (m, 4H), 1.14 – 1.09 (m, 2H). **^13^C NMR** (126 MHz, MeOD) δ 175.58, 171.90, 144.35, 135.97, 126.35, 125.63, 124.47, 59.66, 57.59, 41.92, 39.88, 35.76, 35.64, 35.40, 32.33, 28.17, 25.66, 24.71. **HRMS (ESI):** calcd for C_30_H_37_O_5_N_4_S_2_ [M^+^], *m/z* = 597.21999, found 597.22036. The purity of the compound was analyzed by analytical RP-HPLC. Gradient starts from 5% B for 4 min, 5% - 100% B for 16 min, wash. The product eluted at 18.5 min was detected at λ = 270 nm.

#### Cephalosporin derivative **12b**

RP-HPLC (Gemini-NX): Gradient 5% B for 14 min; 5% - 40% B for 46 min; 40% - 100% B for 5 min, wash. The desired product eluting at 37-38 min was collected and lyophilized to afford product **12b** (4.5 mg, 0.101 mmol, 7%) as a slightly yellowish solid.

**^1^H NMR**(500 MHz, MeOD) δ 9.09 (d, *J* = 6.6 Hz, 2H), 7.95 (d, *J* = 6.6 Hz, 2H), 7.28 – 7.24 (m, 2H), 6.99 – 6.90 (m, 4H), 5.70 (d, *J* = 5.0 Hz, 1H), 5.67 (d, *J* = 14.1 Hz, 1H), 5.19 (d, *J* = 14.1 Hz, 1H), 5.09 – 5.04 (m, 2H), 3.80 (dd, *J* = 11.4, 5.6 Hz, 4H), 3.63 – 3.53 (m, 1H), 3.14 – 3.07 (m, 1H), 2.47 (dq, *J* = 12.4, 6.2 Hz, 1H), 2.35 (t, *J* = 7.4 Hz, 2H), 1.90 (dq, *J* = 12.4, 6.2 Hz, 1H), 1.78 – 1.60 (m, 4H), 1.54 – 1.45 (m, 3H). **HRMS (ESI):** calcd for C_28_H_33_O_5_N_4_S_4_ [M^+^], *m/z* = 633.13283, found 633.13297. The purity of the compound was analyzed by analytical RP-HPLC. Gradient starts from 5% B for 4 min, 5% - 100% B for 16 min, wash. The product eluted at 16.5 min was detected at λ = 270 nm.

#### Cephalosporin derivative **13a**

RP-HPLC (Gemini-NX): Gradient 10% B for 14 min; 10% - 50% B for 36 min; 50% - 100% B for 2 min, wash. The desired product eluting at 26.0–27.5 min was collected and lyophilized to afford product **13a** (6.5 mg, 0.041 mmol, 29%) as a white solid.

**^1^H NMR** (500 MHz, MeOD) δ 9.09 (d, *J* = 6.3 Hz, 2H), 7.92 (d, *J* = 6.3 Hz, 2H), 7.25 (dd, *J* = 5.1, 1.2 Hz, 1H), 6.96 – 6.90 (m, 2H), 5.70 (d, *J* = 4.9 Hz, 1H), 5.67 (d, *J* = 14.0 Hz, 1H), 5.18 (d, *J* = 14.0 Hz, 1H), 5.07 (d, *J* = 4.9 Hz, 1H), 3.82 – 3.75 (m, 2H), 3.59 (d, *J* = 17.8 Hz, 1H), 3.10 (d, *J* = 17.8 Hz, 1H), 2.78 (p, *J* = 8.1 Hz, 1H), 1.98 – 1.88 (m, 2H), 1.80 – 1.70 (m, 4H), 1.68 – 1.60 (m, 2H). **HRMS (ESI)**:calcd for C_26_H_25_O_5_N_4_S_4_ [M^+^], *m/z* = 541.15739, found 541.15767. The purity of the compound was analyzed by analytical RP-HPLC (Gemini-NX). Gradient starts from 5% B for 4 min, 5% - 100% B for 11 min, wash. The product eluted at 15.2 min was detected at λ = 270 nm.

#### Cephalosporin derivative **14**

RP-HPLC (Gemini-NX): Gradient 5% B for 14 min; 5% - 60% B for 51 min; 60% - 100% B for 5 min, wash. The desired product, eluting at 52-53 min, was collected, and lyophilized to afford product **14** (2 mg, 0.045 mmol, 6%) as a yellow solid. **^1^H NMR** (500 MHz, DMSO-*d*6) δ 9.42 (d, *J* = 6.5 Hz, 2H), 9.10 (d, *J* = 5.7 Hz, 1H), 9.07 (d, *J* = 8.3 Hz, 1H), 7.94 (d, *J* = 6.5 Hz, 2H), 7.33 (dd, *J* = 5.1, 1.3 Hz, 1H), 6.92 (dd, *J* = 5.1, 3.4 Hz, 1H), 6.89 – 6.87 (m, 1H), 5.63 (d, *J* = 13.3 Hz, 1H), 5.55 (dd, *J* = 8.3, 4.9 Hz, 1H), 5.04 (d, *J* = 13.7 Hz, 1H), 5.02 (d, *J* = 4.9 Hz, 1H), 4.57 (d, *J* = 5.5 Hz, 1H), 3.70 (d, *J* = 3.0 Hz, 2H), 3.52 (d, *J* = 17.6 Hz, 1H), 3.08 – 2.95 (m, 6H), 2.86 (p, *J* = 7.1 Hz, 1H), 1.46 (s, 18H). ^13^C NMR (126 MHz, DMSO) δ 172.13, 169.92, 167.85, 163.28, 163.02, 162.61, 159.35, 149.42, 144.64, 138.59, 136.97, 126.57, 126.23, 125.25, 124.92, 122.08, 108.50, 85.24, 60.93, 58.98, 57.35, 45.82, 41.82, 35.76, 31.89, 27.80, 24.31. **HRMS (ESI)**: calcd for C_34_H_43_O_9_N_4_S_4_ [M+H]^+^, *m/z* = 779.19074, found 779.19080. The purity of the compound was analyzed by analytical RP-HPLC (Gemini). Gradient starts from 5% B for 4 min, 5% - 100% B for 11 min, wash. The product eluted at 16.2 min was detected at λ = 270 nm.

##### Synthesis of cephalosporin derivative 13b

Cephalosporin derivative **14** (4.7 mg, 0.006 mmol) was dissolved in DCM (0.450 mL) and TFA (0.140 mL) was added. The reaction was stirred at rt for 1 h. Then the solvent was evaporated and the crude product was washed with IPE (2×1 mL). The crude product was dissolved in DMSO (0.200 mL) and the reaction mixture was stirred at 70 °C for 4 h in open flask. Methanol (0.200 mL) was added into the reaction. The mixture was filtered through the syringe filter (1.0 μm) and purified by semi-prep RP-HPLC (Hydro): gradient 5% B for 7 min; 5% - 30% B for 28 min; 30% - 100% B for 2 min, wash. The desired product **13b**, eluting at 25-30 min, was collected, and lyophilized to afford product as a yellowish solid (2.1 mg, 0.006 mmol, 60%). **^1^H NMR** (500 MHz, DMSO-*d*_6_) δ 9.41 (d, *J* = 6.2 Hz, 2H), 9.10 – 9.00 (m, 2H), 7.97 (d, *J* = 6.1 Hz, 2H), 7.34 (d, *J* = 4.9 Hz, 1H), 6.92 (dd, *J* = 3.5, 4.9 Hz, 1H), 6.90 – 6.87 (m, 1H), 5.64 (d, *J* = 13.3 Hz, 1H), 5.54 (dd, *J* = 8.3, 4.8 Hz, 1H), 5.01 (q, *J* = 4.2 Hz, 2H), 4.58 (d, *J* = 5.6 Hz, 2H), 3.71 (d, *J* = 1.9 Hz, 2H), 3.52 (d, *J* = 17.6 Hz, 2H), 3.49 – 3.43 (m, 3H), 3.00 (d, *J* = 17.6 Hz, 1H), 1.03 (d, *J* = 6.1 Hz, 1H). **HRMS (ESI)**:calcd for C_24_H_25_O_5_N_4_S_4_ [M]^+^, *m/z* = 577.07023, found 577.07039. The purity of the compound was analyzed by analytical RP-HPLC (Gemini-NX). Gradient starts from 5% B for 4 min, 5% - 100% B for 11 min, wash. The product eluted at 15.2 min was detected at λ = 270 nm.

#### Synthesis of cephalosporin derivative 11

Under an argon atmosphere, a mixture of cephalothin **3** (46 mg, 0.115 mmol) and MSTFA (0.040 mL, 0.216 mmol) in dry DCM (0.700 mL) in a Schlenk tube, was stirred at rt for 15 min until the cephalothin was dissolved. Then iodotrimethylsilane (0.020 mL, 0.144 mmol) was added, and stirring was continued at rt for 1 h. The mixture was concentrated under reduced pressure connected to Schlenk line, and the residual oil was dissolved in dry DMF (0.700 mL). A few drops of dry THF were added, and the mixture was stirred for 1 minute. Tert-butyl pyridin-4-ylmethylcarbamate (15 mg, 0.072 mmol) dissolved in dry DMF (0. 700 mL) was added and the reaction mixture was stirred at rt for 1 h. The solvents were evaporated, the crude was resuspended in DCM (0.500 mL) and TFA (0.200 mL) was added. The mixture was stirred at rt for 1 h. The solvent was evaporated and the mixture was dissolved in water/MeOH. The formed precipitate was filtered out and the solution was purified by preparative RP-HPLC (Hydro): gradient 5% B for 14 min; 5% - 15% B for 16 min; 15% - 100% B for 3 min, wash. The desired product **11**, eluting at 9-11 min, was collected and lyophilized to afford product **11** as a white solid (18 mg, 0.072 mmol, 56%). **^1^H NMR** (500 MHz, MeOD) δ 9.21 (d, *J* = 6.6 Hz, 2H), 8.16 (d, *J* = 6.6 Hz, 2H), 7.25 (dd, *J* = 5.1, 1.3 Hz, 1H), 6.97 – 6.91 (m, 2H), 5.75 (d, *J* = 14.2 Hz, 1H), 5.70 (d, *J* = 5.0 Hz, 1H), 5.32 (d, *J* = 14.2 Hz, 1H), 5.12 (d, *J* = 5.0 Hz, 1H), 4.53 (s, 1H), 3.83 – 3.75 (m, 2H), 3.66 (d, *J* = 18.1 Hz, 1H), 3.20 (d, *J* = 18.1 Hz, 1H). **^13^C NMR**(126 MHz, MeOD) δ 173.32, 146.44, 137.31, 128.46, 127.81, 127.79, 125.94, 61.28, 59.10, 42.64, 37.13. **HRMS (ESI):** calcd for C_20_H_21_O_4_N_4_S_2_ [M^+^], *m/z* = 445.09987, found 445.09929. The purity of the compound was analyzed by analytical RP-HPLC (Gemini-NX). Gradient starts from 5% B for 4 min, 5% - 100% B for 11 min, wash. The product eluted at 12.2 min was detected at λ= 270 nm.

### Procedure for the synthesis of cephalosporin derivatives with amino thiadiazol side chain

#### General procedure

Under an argon atmosphere, NaI (3 equiv.) was added to a mixture of cephalosporin derivative **15** (1.5 equiv.) in dry acetone (0.1 M solution). The reaction mixture was stirred at rt for 40 min.Subsequently, the pyridine derivative (1 equiv.) in dry acetone (0.1 M solution) was added and the reaction was stirred at rt for 2 – 4 h. After the reaction was finished, the solvent was evaporated and the mixture was washed with IPE (2 mL). The formed precipitate was filtered and then dissolved in mixture of DCM/anisole/TFA 5:1:1 (0.1 mL). The reaction mixture was stirred for 1-2 h and then IPE (2 mL)was added to the mixture. The resulting suspension was centrifuged. The supernatant was removed, and the precipitate was washed with IPE two more times. The crude was dissolved in mixture of MeOH/H_2_O/CH_3_CN, filtered through a SPE column, and purified by preparative RP-HPLC. The purification methods, analytical data, and yields are below.

#### Cephalosporin derivative **17a**

RP-HPLC (Gemini-NX): Gradient 5% B for 14 min; 5% - 45% B for 46 min; 45% - 100% B for 5 min, wash. The desired product, eluting at 41 min, was collected and lyophilized to afford product **17a** (9.2 mg, 0.115 mmol, 12%) as a white solid. **^1^H NMR** (500 MHz, MeOD) δ 9.09 (d, *J* = 6.3 Hz, 2H), 8.49 – 8.44 (m, 1H), 7.93 (d, *J* = 6.3 Hz, 2H), 5.86 (d, *J* = 4.9 Hz, 1H), 5.67 (d, *J* = 13.8 Hz, 1H), 5.19 (d, *J* = 13.8 Hz, 1H), 5.15 (d, *J* = 4.9 Hz, 1H), 4.01 (s, 3H), 3.60 (d, *J* = 17.8 Hz, 2H), 3.09 (d, *J* = 17.8 Hz, 1H), 2.33 (t, *J* = 7.6 Hz, 1H), 2.36-2.25 (m, 1H), 1.81 – 1.73 (m, 4H), 1.68 – 1.59 (m, 6H), 1.55 – 1.50 (m, 2H), 1.38 – 1.33 (m, 3H), 1.17 – 1.04 (m, 3H). **HRMS (ESI):** calcd for C_29_H_37_O_6_N_8_S_2_ [M^+^], *m/z* = 657.22720, found 657.22748. The purity of the compound was analyzed by analytical RP-HPLC (Gemini-NX). Gradient starts from 5% B for 4 min, 5% - 100% B for 11 min, wash. The product eluted at 14.7 min was detected at λ=270 nm.

#### Cephalosporin derivative **17b**

RP-HPLC: Gradient 10% B for 14 min; 10% - 40% B for 46 min; 45% - 100% B for 5 min, wash. The desired product, eluting at 33.8 min, was collected and lyophilized to afford product **17b** (2.56 mg, 0.067 mmol, 6%) as a white solid. **^1^H NMR** (500 MHz, MeOD) δ 9.11 (d, *J* = 6.3 Hz, 2H), 8.92 – 8.88 (m, 1H), 7.95 (d, *J* = 6.3 Hz, 2H), 5.79 (d, *J* = 5.0 Hz, 1H), 5.69 (d, *J* = 15.3 Hz, 1H), 5.18 (d, *J* = 14.0 Hz, 1H), 5.14 (d, *J* = 4.9 Hz, 1H), 4.04 (s, 3H), 3.21 – 3.15 (m, 2H), 3.15 – 3.09 (m, 2H), 2.50 – 2.41 (m, 2H), 2.36 (t, *J* = 7.4 Hz, 2H), 2.34 – 2.28 (m, 1H), 1.94 – 1.86 (m, 2H), 1.77 – 1.60 (m, 6H), 1.56 – 1.44 (m, 4H). **HRMS (ESI)**:calcd for C_27_H_33_O_6_N_9_S_4_ [M]^+^, *m/z* = 693.14004, found 693.13968. The purity of the compound was analyzed by analytical RP-HPLC (Gemini-NX). Gradient starts from 5% B for 4 min, 5% - 100% B for 11 min, wash. The product eluted at 15.3 min was detected at λ= 270 nm.

#### Cephalosporin derivative **18a.**

RP-HPLC (Gemini-NX): Gradient 5% B for 14 min; 5% - 30% B for 46 min; 30% - 100% B for 5 min, wash. The desired product, eluting at 25.5 min, was collected and lyophilized to afford product **18a** (4.5 mg, 0.049 mmol, 15%) as a white solid. **^1^H NMR** (500 MHz, DMSO-*d*6) δ 9.46 (d, *J* = 8.3 Hz, 1H), 9.38 (d, *J* = 6.3 Hz, 2H), 8.61 −8.59 (m, 1H), 8.12 – 8.08 (m, 3H), 7.91 (d, *J* = 6.3 Hz, 2H), 5.65 (dd, *J* = 8.5, 5.0 Hz, 1H), 5.62 (d, *J* = 13.5 Hz, 1H), 5.05 (d, *J* = 5.0 Hz, 1H), 5.02 (d, *J* = 14.1 Hz, 1H), 4.50 (d, *J* = 5.7 Hz, 2H), 3.85 (s, 3H), 3.50 (d, *J* = 17.5 Hz, 1H), 2.98 (d, *J* = 17.5 Hz, 1H), 2.72 – 2.64 (m, 1H), 1.85 – 1.78 (m, 2H), 1.69 – 1.59 (m, 4H), 1.54 – 1.49 (m, 2H). **HRMS (ESI):** calcd for C_25_H_29_O_5_N_4_S_2_ [M^+^], *m/z* = 601.16460, found 601.16508. The purity of the compound was analyzed by analytical RF-HPLC (Gemini). Gradient starts from 5% B for 4 min, 5% - 100% B for 11 min, wash. The product eluted at 12.1 min was detected at λ = 270 nm.

#### Cephalosporin derivative **18b.**

Semi-prep RP-HPLC (Hydro): Gradient 5% B for 7 min; 5% - 30% B for 23 min; 30% - 100% B for 2 min, wash. The desired product, eluting at 17.5 min, was collected and lyophilized to afford product **18b** (1.2 mg, 0.083 mmol, 2%) as a yellowish solid.

**^1^H NMR** (500 MHz, MeOD) δ 9.10 (d, *J* = 6.5 Hz, 2H), 7.95 (d, *J* = 6.5 Hz, 2H), 5.85 (d, *J* = 4.9 Hz, 1H), 5.69 (d, *J* = 13.5 Hz, 1H), 5.20 (s, 1H), 5.15 (d, *J* = 5.0 Hz, 1H), 4.66 (s, 2H), 4.01 (s, 3H), 3.69 – 3.63 (m, 2H), 3.61 (d, *J* = 17.8 Hz, 1H), 3.45 – 3.38 (m, 5H), 3.10 (d, *J* = 17.8 Hz, 1H). **HRMS (ESI):** calcd for C23H25O6N8S4 [M^+^], *m/z* = 637.07744, found 637.07765. The purity of the compound was analyzed by analytical RP-HPLC (Gemini-NX): Gradient 5% B for 4 min, 5% - 100% B for 11 min, wash. The product eluted at 13.8 min was detected at λ = 270 nm.

### Biology

#### Bacterial strains and growth conditions

*Bacillus subtilis* (*B. subtilis* ATCC 6633), *Staphylococcus aureus (S. aureus* ATCC 25922, ATCC 29213, ATCC 43300 (MRSA)), *Enterococcus faecalis (E. faecalis* ATCC 51299 (VanB)), *Escherichia coli (E. coli* ATCC 25922, K12 MG1622), *Pseudomonas aeruginosa (P. aeruginosa* ATCC 27853, PAO1) was purchased from either the German Collection of Microorganisms and Cell Cultures (DSMZ) or the American Type Culture Collection (ATCC). The bacterial culture was stored at −80 °C, and new cultures were prepared by streaking on Mueller Hinton (MH) Agar, Luria-Bertani (LB) Agar, Trypsic Soy (TS) Agar plates. The overnight culture was prepared by inoculating a single colony into a sterile tube (15 mL) containing the MH, LB, or TS broth (5 mL) and the cultures were shaken (200 rcf/min) overnight at 37 °C.

#### Determination of Minimum Inhibitory Concentration (MIC)

The minimum inhibitory concentration (MIC) was determined using the broth microdilution method according to the guidelines outlined by the European Committee on Antimicrobial Susceptibility Testing (EUCAST) standard protocol.^25^ See the Supporting Information for the detailed procedure.

#### Defining conditions for biofilm inhibition assay

##### Biofilm production

The conditions for biofilm formation of *S. aureus* ATCC 29213, *S. aureus* ATCC 43300, and *E. faecalis* ATCC 51299 were published earlier.^25^ The overnight culture of *E. coli* (ATCC 25922, K12 MG1622) and *P. aeruginosa* (ATCC 27853, PAO1) was diluted in LB broth to OD_600_ = 0.1 – 0.12 corresponding to 10 ×10^8^ cells/mL. The broths used for the experiment were prepared: Muller-Hilton broth (MHB), Trypsic Soy (TS), Trypsic Soy supplemented with 1% glucose (TSG), Trypsic Soy supplemented with 2% glucose (TS2G),

Brain Heart Infusion (BHI), BHI supplemented with 1% glucose (BHIG), M63 minimal supplemented with MgSO_4_, glucose and casamino acids (M63), M63 minimal supplemented with arginine (M63A). The adjusted to OD600 cells were diluted 1:100 with corresponding medium. An aliquot of 100 ul of bacterial suspension per well was dispensed into a 96-well round bottom microplate. Microplates were then incubated at 37 °C for 24 h.

##### Assessment of biofilm biomass by crystal violet

An aliquot of 0.130 ul of 0.1% solution of crystal violet in water was added in each well contained biofilm. The microplates were incubated for 10 min at rt and the plates were rinsed 2-3 times with sterile water. The plates were left to dry for 2 h at rt. Next, 0.130 ul of 30% AcOH in water was added to each well of the microplate to solubilize the CV. The plates were incubated for 10 min at rt. The obtained solutions were transferred into a new flat bottom 96-well plate. The absorbance was quantified at 570 nm using 30% acetic acid in water as the blank by a plate reader (Synergy H1 from BioTek).

##### Assessment of metabolic activity of biofilm cells by fluorescein diacetate (FDA)

Biofilm production was performed using M63A for *P. aeruginosa* ATCC 27853. A stock of FDA (Acros) was prepared at 1 mg/mL in DMSO, respectively. The solutions were filter-sterilized and stored at 4 °C in the dark. Three concentrations of FDA solution were investigated: 2 μg/mL, 4 μg/mL and 8 μg/mL at 37 °C. The diluted FDA solutions in PBS were prepared right before the assay. For the assay, firstly, biofilm was carefully washed with 200 μL of sterile water. Next, 100 μL of diluted FDA solutions were added into each well containing biofilm, along with its respective negative controls (un-inoculated broth, three wells). Microplates were placed in a plate reader to measure the kinetics of the reaction. The relative fluorescence units (RFU) (λ_Ex_=490 nm and λ_Em_=526 nm) were measured every 2 min for 2 h (Figure S1, Supporting Information). The experiment was performed twice in triplicates. The optimal conditions were chosen based on the analysis of quality parameters (Z’□>□0.50) calculated by the equations (Table S3, Supporting Information).

#### Determination of Minimum Biofilm Reduction Concentration (MBRC)

Cephalosporin derivatives stock solutions were prepared in sterile water or water + 5% DMSO to a concentration of 1 mg/mL. MBRC assay was performed by the broth microdilution method in 96-well round bottom microplate adapted from Clinical and Laboratory Standards Institute (CLSI) guideline.^47^

Bacterial suspension was diluted with BHIG broth to obtain an inoculum of 1×10^6^ CFU/mL. Two-fold serial dilutions of the compounds (ranging from 64 μg/ml to 0.06 μg/ml for cephalosporin derivatives) were prepared in BHIG (for *S. aureus* and *E. faecalis)* and M63A (for *P. aeruginosa)* to a final volume of 50 μl. The bacterial suspension (50 μl) was added into each well on the microtiter plate for inoculation, corresponding to approximately 5×10^5^ CFU/ml. The plates were incubated without shaking for 24 h at 37 °C, and MBIC was defined by performing the optimized resazurin or FDA assays (Table S4, Supporting Information). CFUs were also determined after resazurin or FDA assays completion, in order to compare the results. The experiments were performed in triplicates.

#### Statistical analysis

The MIC and MBRC experiments were performed in triplicates. Assessment of the metabolic activity of biofilm cells by fluorescein diacetate (FDA) was performed from two independent experiments, each in triplicates. Statistical quality parameters were calculated according to the methods described before.^48,49^ GraphPad Prism 9 was used for statistical analysis and representation. Results were considered statistically significant when *p* ≤ 0.05. The formulae used for data viability calculation are given in the Supporting Information.

## Data availability

All data generated or analyzed during this study are included in this published article and in the Supplementary Information files.

## Supporting Information

Supporting Information Available: NMR data, detailed experimental procedure and results for MIC, detailed MBRC results, assessment of metabolic activity of biofilm cells and calculation of data viability.

## Acknowledgements

The authors acknowledge the Swiss National Science Foundation (SNSF, Grant 182043) and a Bundesstipendium for financial support. We thank Laura Kqiku for initial experiments not included in this study. The authors acknowledge the NMR and mass spectrometry facilities at the University of Zurich for training and maintenance of the instruments.

## Author Contributions

I.S.S. and K.G. designed the study, I.S.S. performed all experiments, I.S.S. and K.G. analyzed the data, and I.S.S. and K.G. wrote the manuscript.

## Competing interests

The authors declare no competing interests.

## Notes

### Competing Interest Statement

The authors have declared no competing interest.

### Summary of Updates

Update manuscript based on peer review

