## Supplementary material for "Synthesis and antimicrobial evaluation of new cephalosporin derivatives containing cyclic disulfide moieties": SI

**Table of contents**

Page S2: Materials and instrumentation (I)

Page S3: Characterization of all the novel compounds (NMR spectra and HPLC chromatograms) (II)

Page S24: Microbiological assays (III)

I. **Materials and instrumentation**

**Analytical thin-layer chromatography** (TLC) was run on Merck TLC plates silica gel 60 F254 on glass plates with the indicated solvent system; the spots were visualized by UV light (365 nm), and stained by anisaldehyde, ninhydrin, or KMnO_4_ stain. Silica gel **column chromatography** was performed using silica gel 60 (230−400 Mesh) purchased from Sigma-Aldrich with the solvent mixture indicated. The SPE columns used were DSC-18 (Supelco, Sigma). **High performance liquid chromatography** (HPLC) were performed on a Shimadzu HPLC system (LC-20AP dual pump, CBM-20A Communication Bus Module, SPP-20, A UV/vis Detector, FRC-10A Fraction collector) using reverse-phase (RP) columns analytical: Gemini-NX C18 (150 mm x 4.6 mm; 3 µm, 10 Å), Synergy Hydro-RP (150 mm x 4.6 mm; 4 µm, 80 Å), with a flow of 0.4 mL/min; preparative: Gemini-NX C18 (250 mm × 21.2 mm; 10 μm, 110 Å) or Synergy Hydro-RP (250 mm × 21.2 mm; 10 μm, 80 Å) with a flow of 20 mL/min (unless otherwise stated). A solvent system composed of A (H_2_O + 0.1% HCO_2_H) and B (MeCN + 0.1% HCO_2_H). **Ultrahigh-performance liquid chromatography** (UHPLC) coupled to mass spectrometer (MS) experiments were performed on an Ultimate 3000 LC system (HPG-3400 RS pump, WPS-3000 TRS autosampler, TCC-3000 RS column oven, Vanquish DAD detector from Thermo Scientific) coupled to a triple quadrupole (TSQ Quantum Ultra from Thermo Scientific). The separation was performed using a RP column (Kinetex EVO C18; 50 × 2.1 mm; 1.7 μm; 100 Å, Phenomenex), a flow of 0.4 mL/min, a solvent system composed of A (H_2_O + 0.1% HCO_2_H) and B (MeCN + 0.1% HCO_2_H) and an elution gradient starting with 5% B, increasing from 5−95% B in 3.5 min, from 95−100% B in 0.05 min, and washing the column with 100% B for 1.25 min. The filtration before UHPLC analysis were performed using Syringe Filters, Nylon 66, 0.22 µm. **IR-spectroscopy** was performed on a Varian 800 FT-IR ATR Spectrometer. **Lyophilization** was performed on a Christ Freeze-dryer ALPHA 1−4 LD plus. **High-resolution electrospray mass spectra** (HRMS (ESI)) were recorded on a timsTOF Pro TIMS-QTOF-MS instrument (Bruker Daltonics GmbH, Bremen, Germany). The samples were dissolved in (e.g., MeOH) at a concentration of ca. 50 μg/mL and analyzed via continuous flow injection (2 μL/ min). The mass spectrometer was operated in the positive (or negative) electrospray ionization mode at 4000 V (−4000 V) capillary voltage and −500 V (500 V) end plate offset with a N_2_ nebulizer pressure of 0.4 bar and a dry gas flow of 4 mL/min at 180 °C. Mass spectra were acquired in a mass range from m/ z 50 to 2000 at ca. 20 000 resolution (m/z 622) and at 1.0 Hz rate. The mass analyzer was calibrated between m/z 118 and 2721 using an Agilent ESI-L low concentration tuning mix solution (Agilent, USA) at a resolution of 20 000 giving a mass accuracy below 2 ppm. All solvents used were purchased in best LC-MS quality.

II. **Characterization of all the novel compounds (NMR spectra and HPLC chromatograms)**

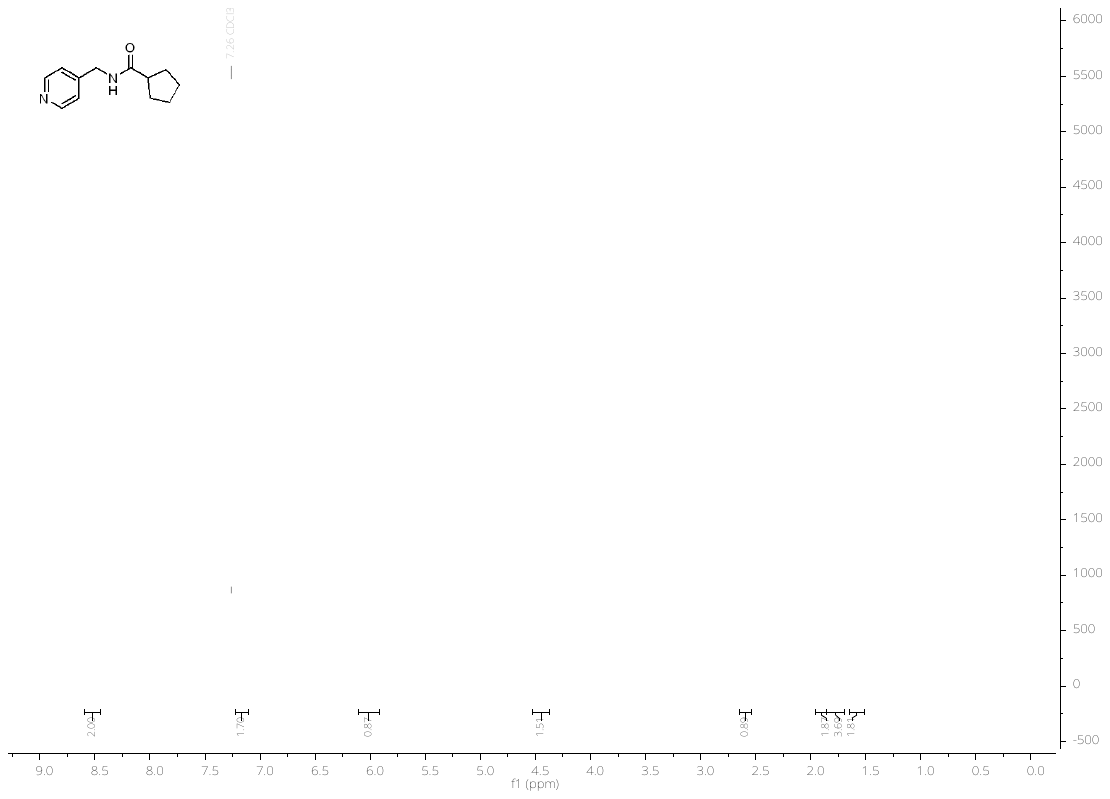

500 MHz ^1^H-NMR in CDCl_3_

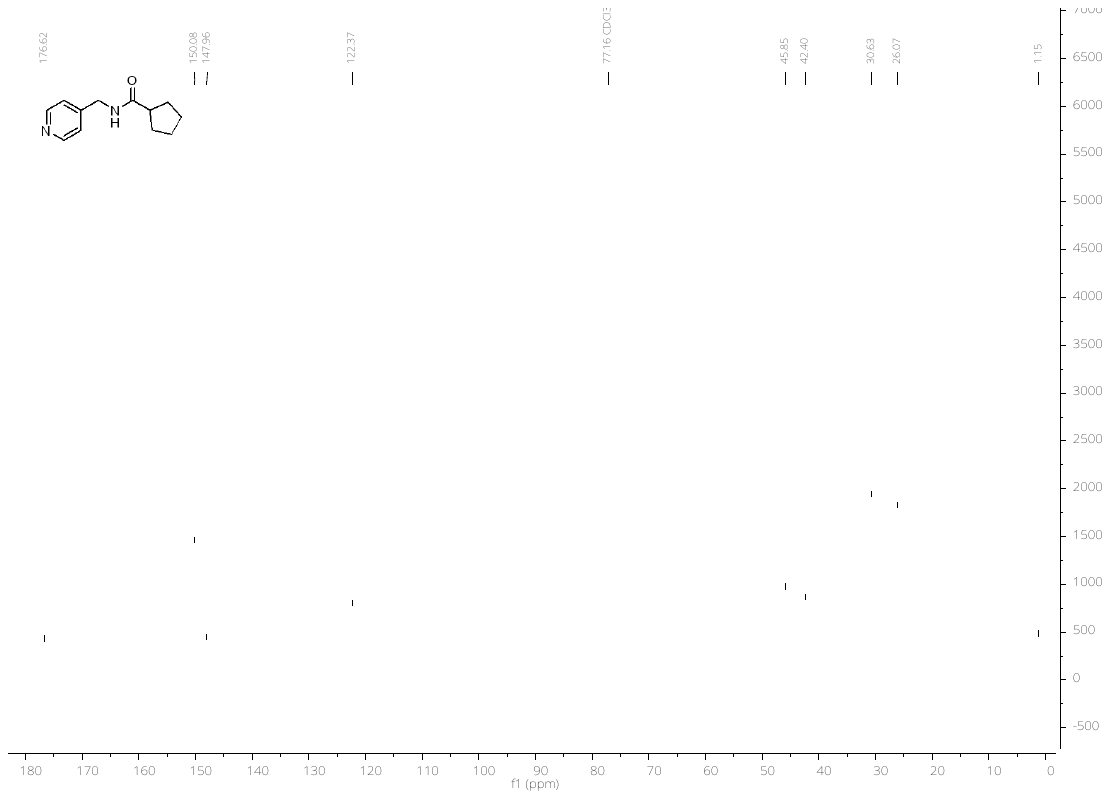

126 MHz ^13^C-NMR in CDCl_3_

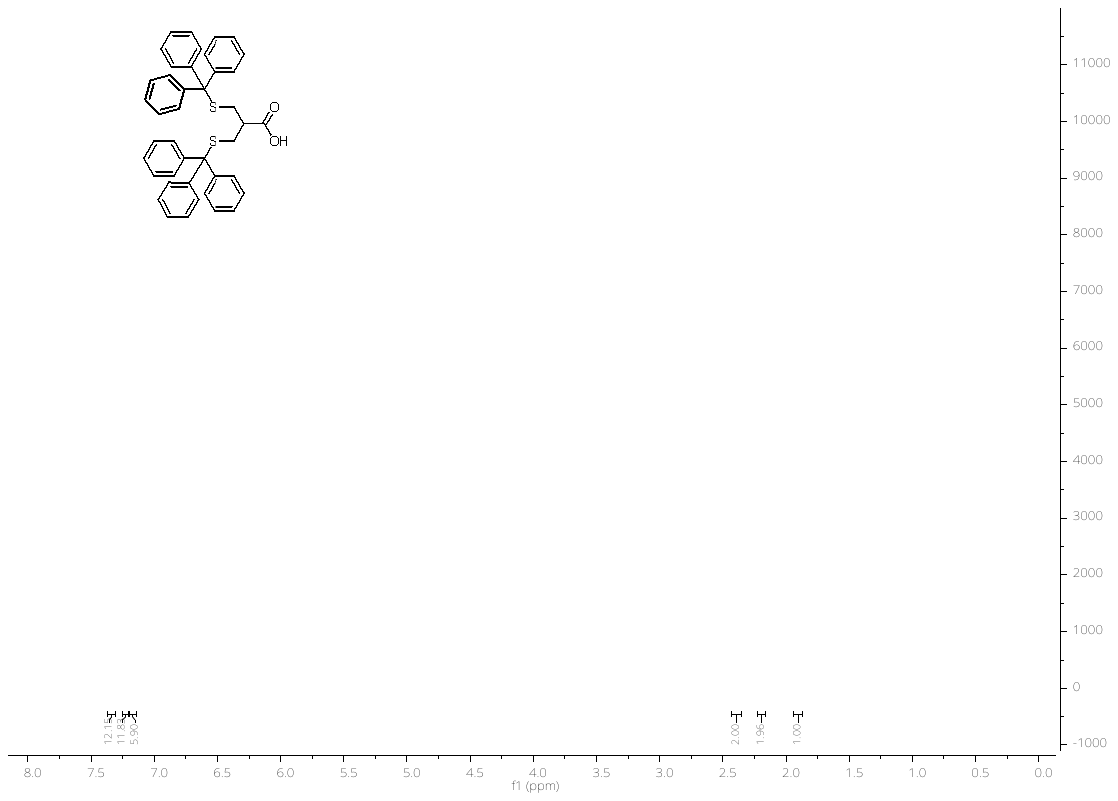

500 MHz ^1^H-NMR in CDCl_3_

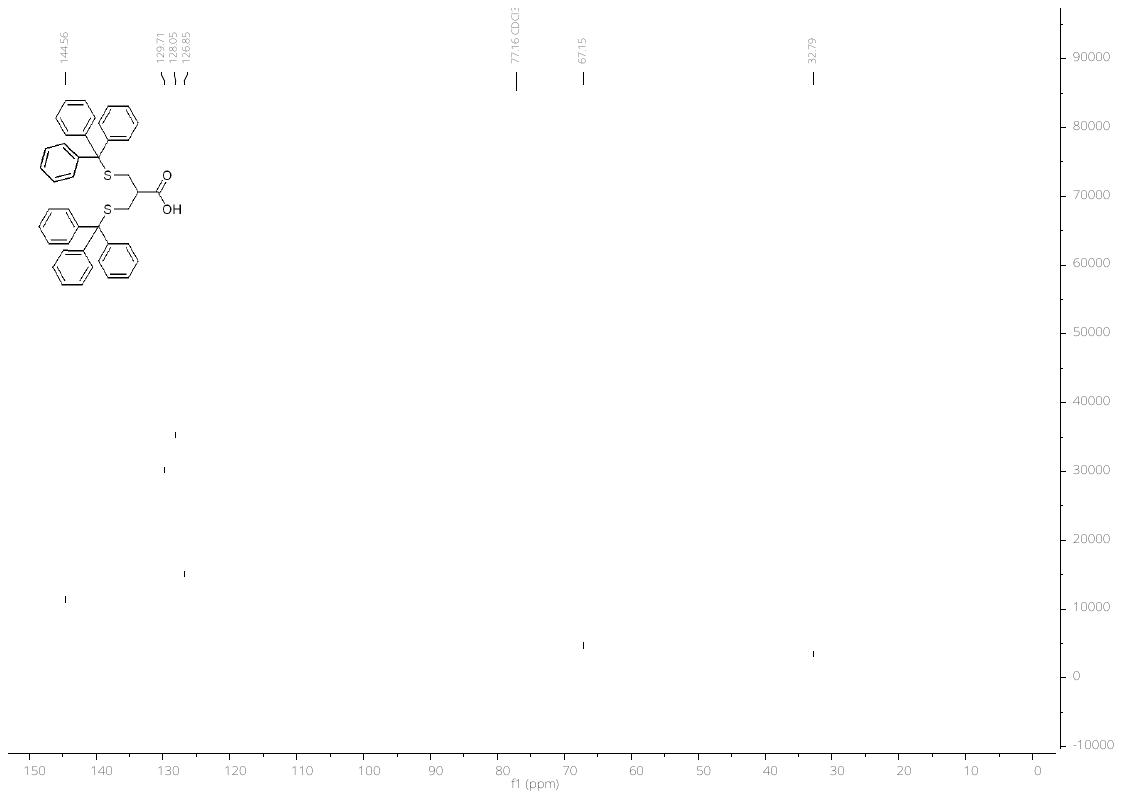

126 MHz ^13^C-NMR in CDCl_3_

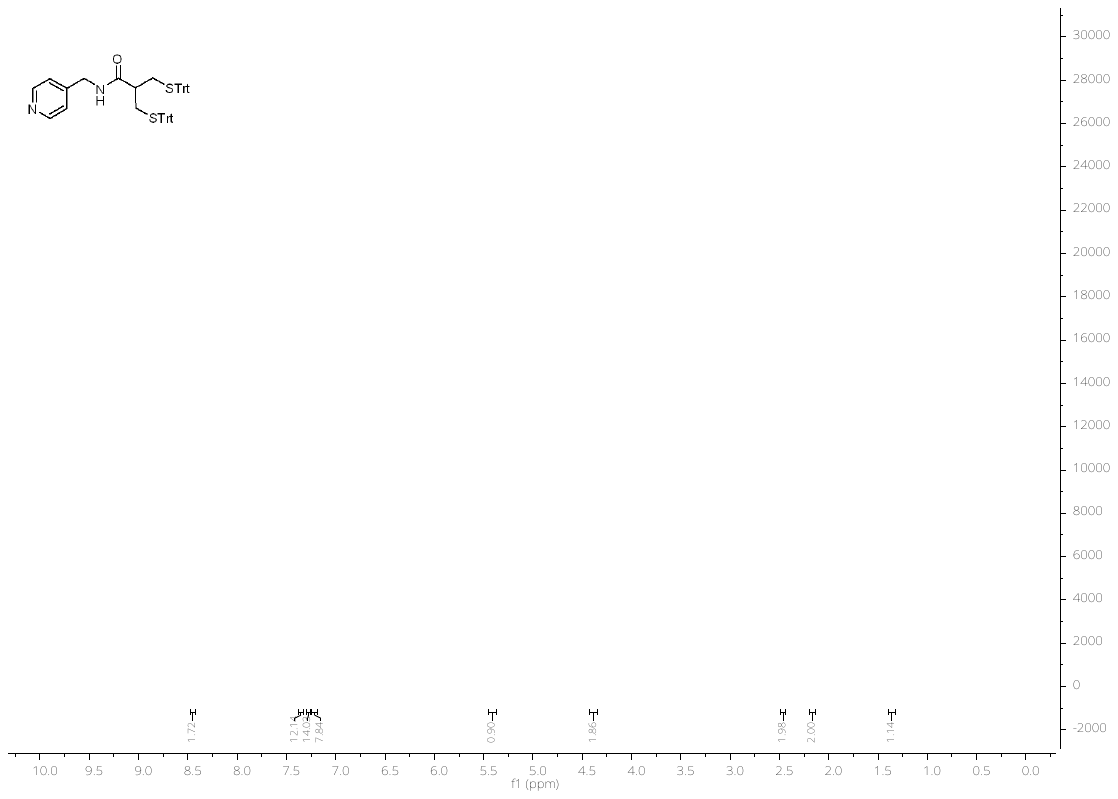

500 MHz ^1^H-NMR in CDCl_3_

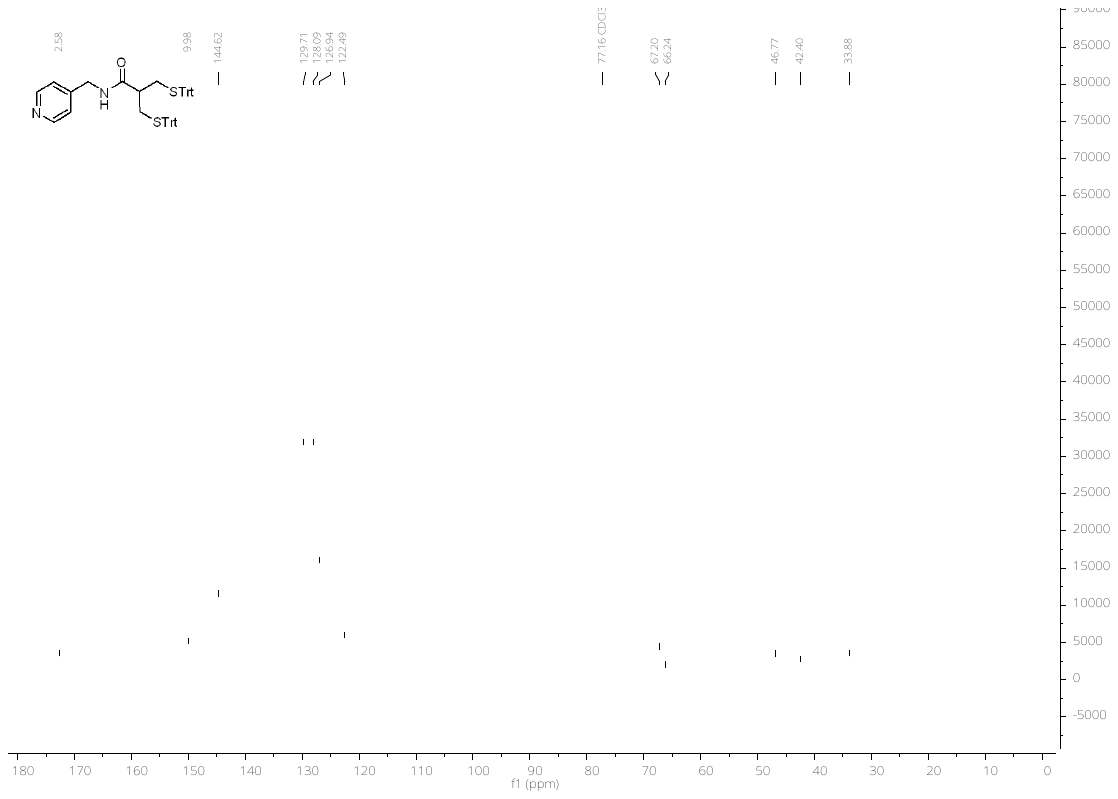

126 MHz ^13^C-NMR in CDCl_3_

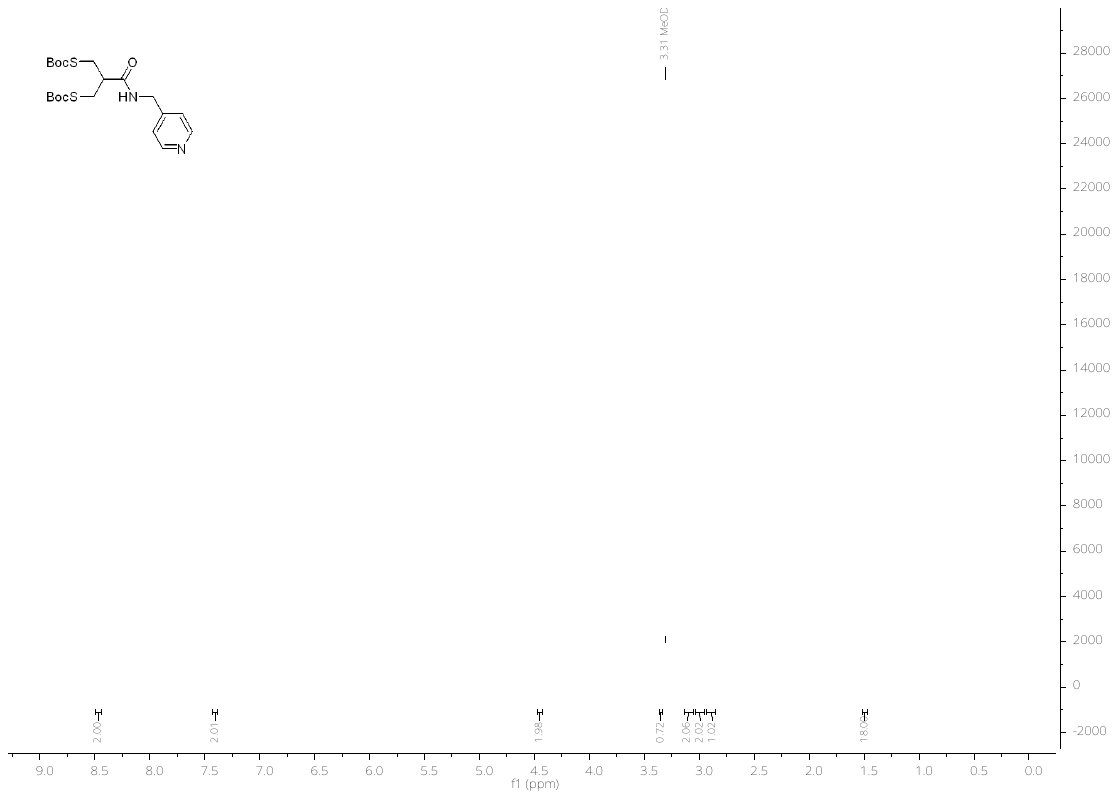

400 MHz ^1^H-NMR in MeOD

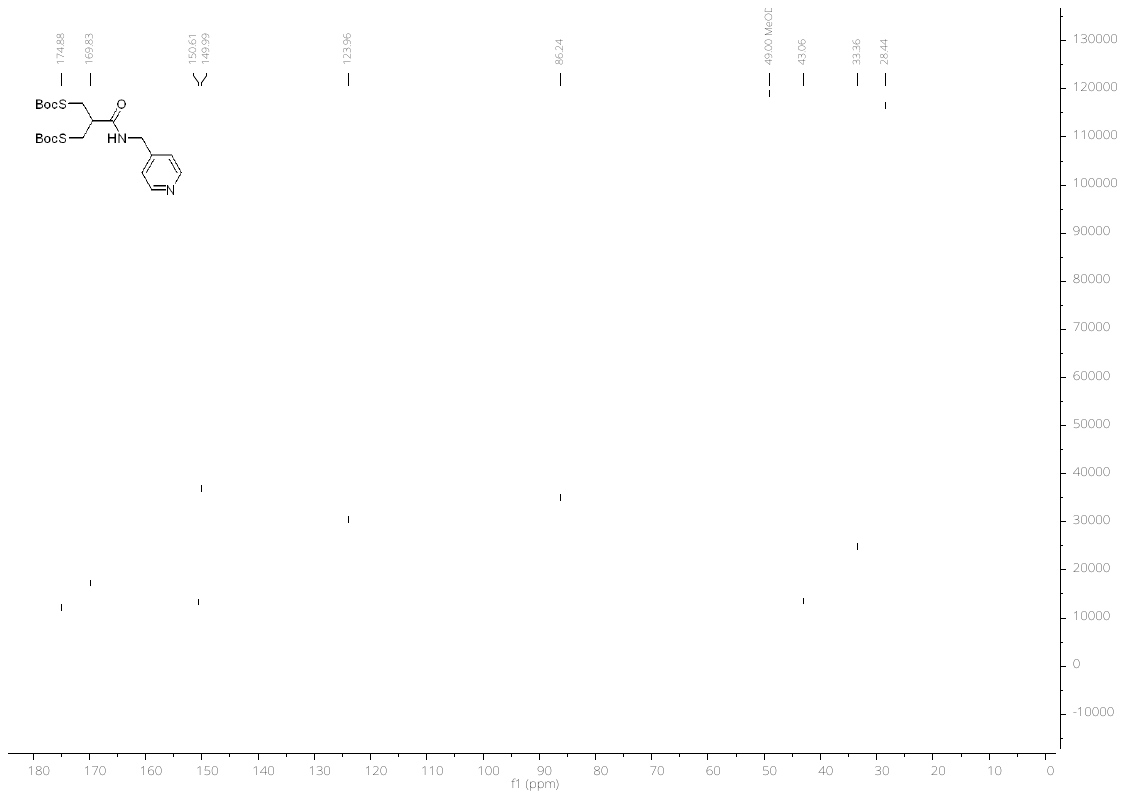

101 MHz ^13^C-NMR in MeOD

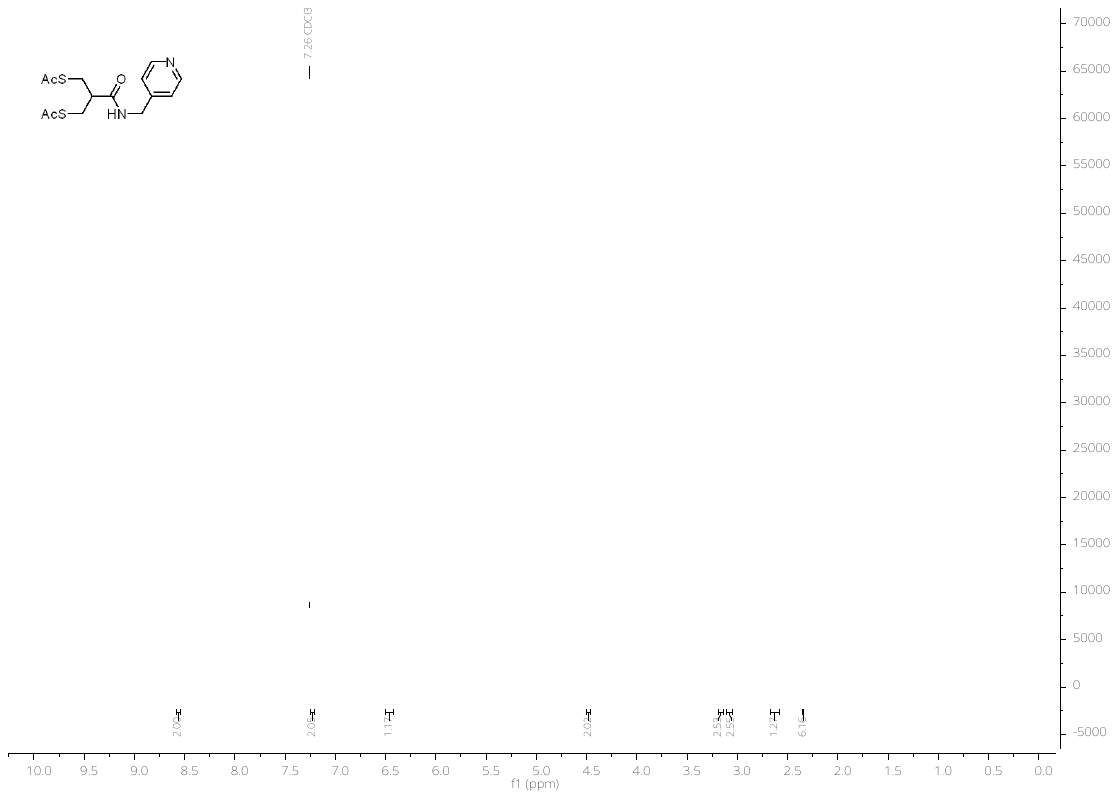

500 MHz ^1^H-NMR in CDCl_3_

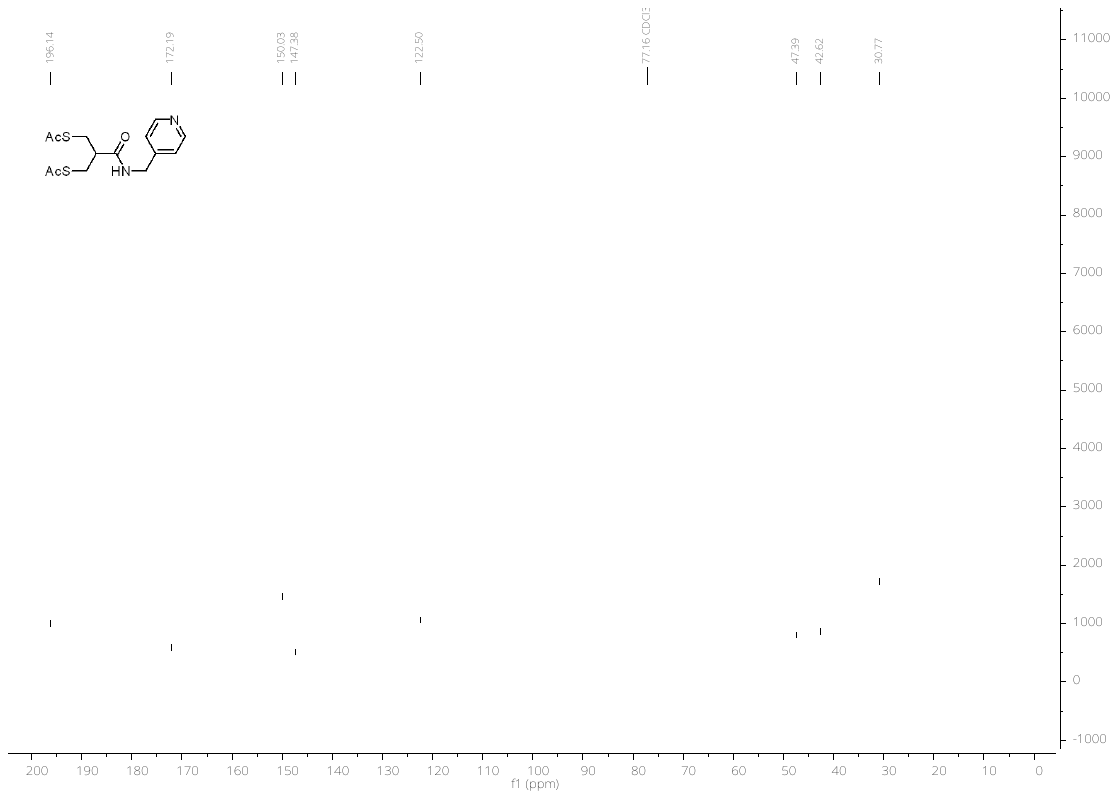

126 MHz ^13^C-NMR in CDCl_3_

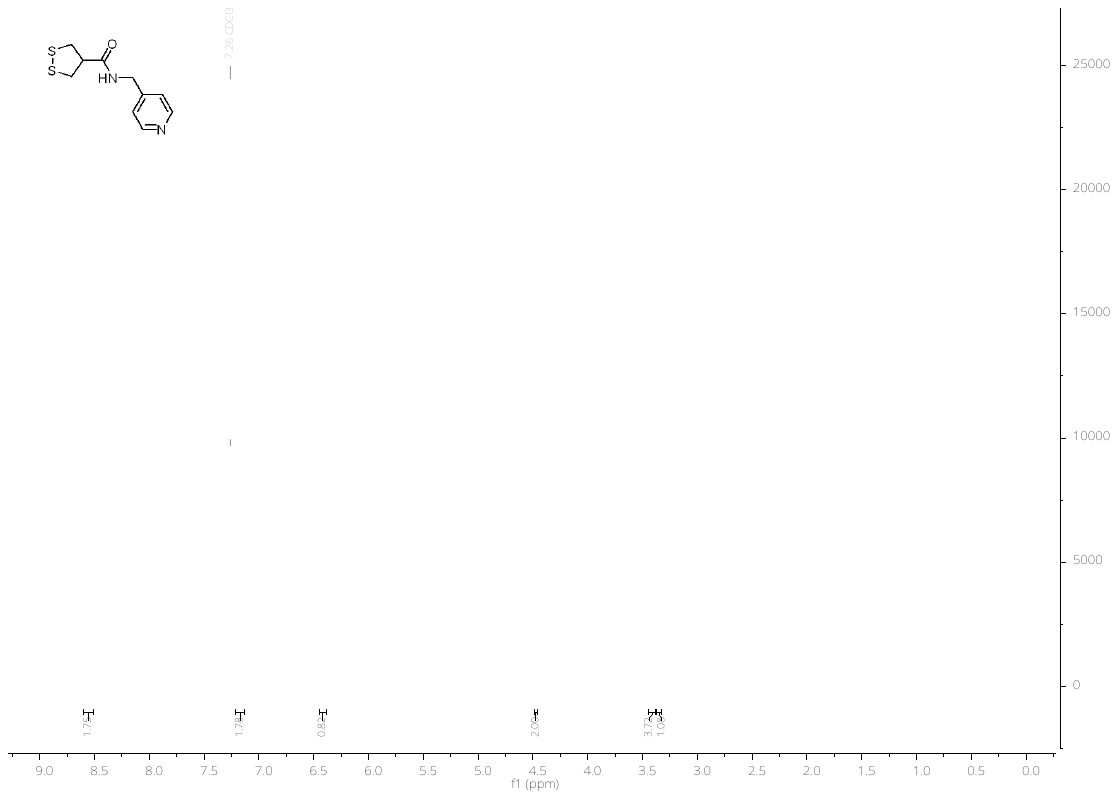

500 MHz ^1^H-NMR in CDCl_3_

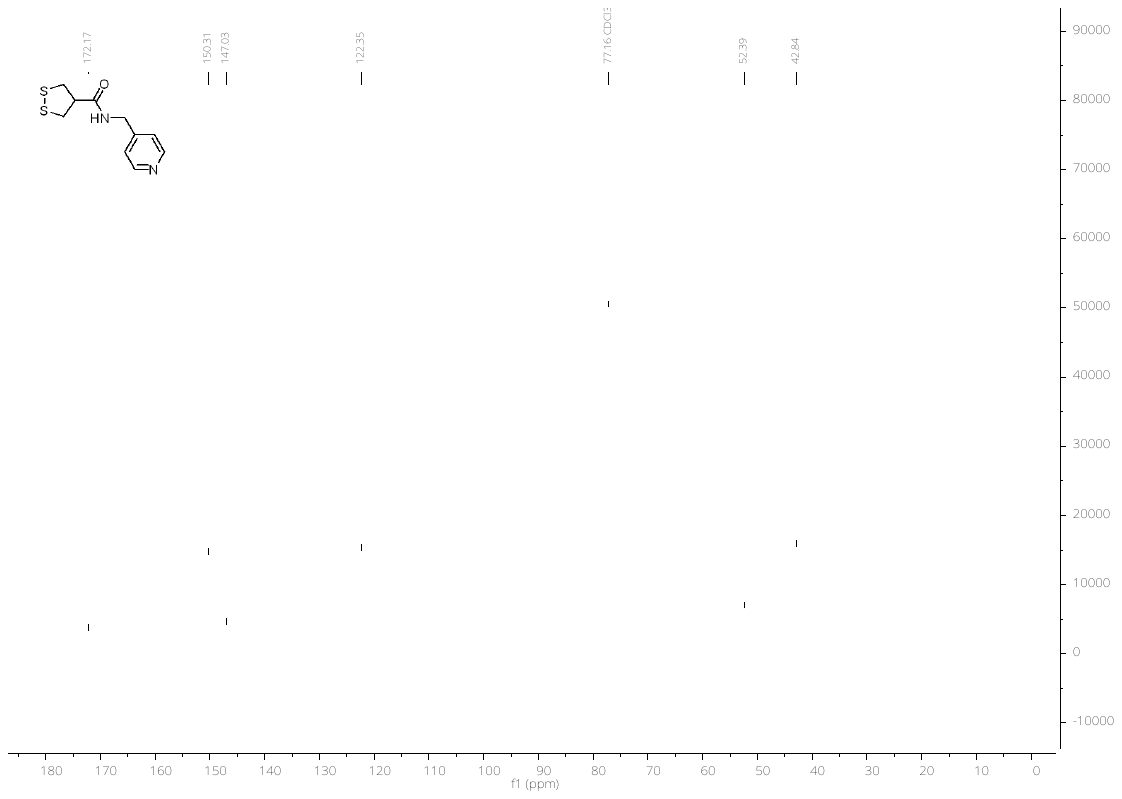

126 MHz ^13^C-NMR in CDCl_3_

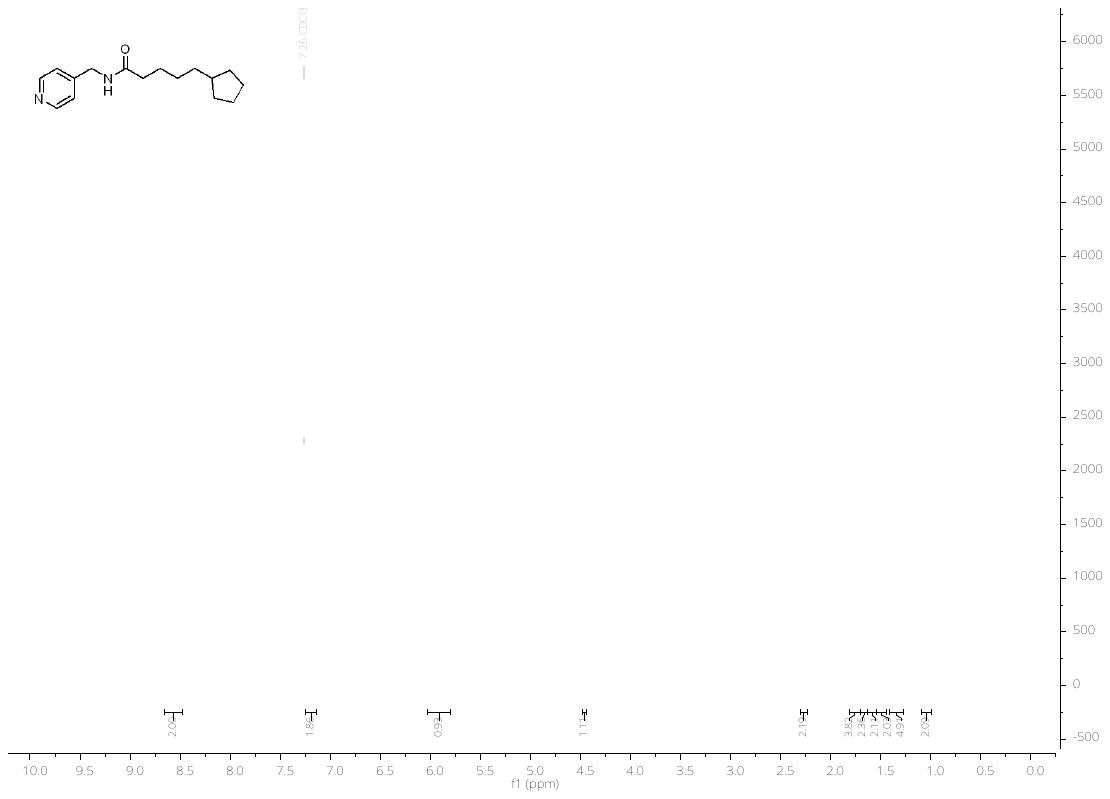

500 MHz ^1^H-NMR in CDCl_3_

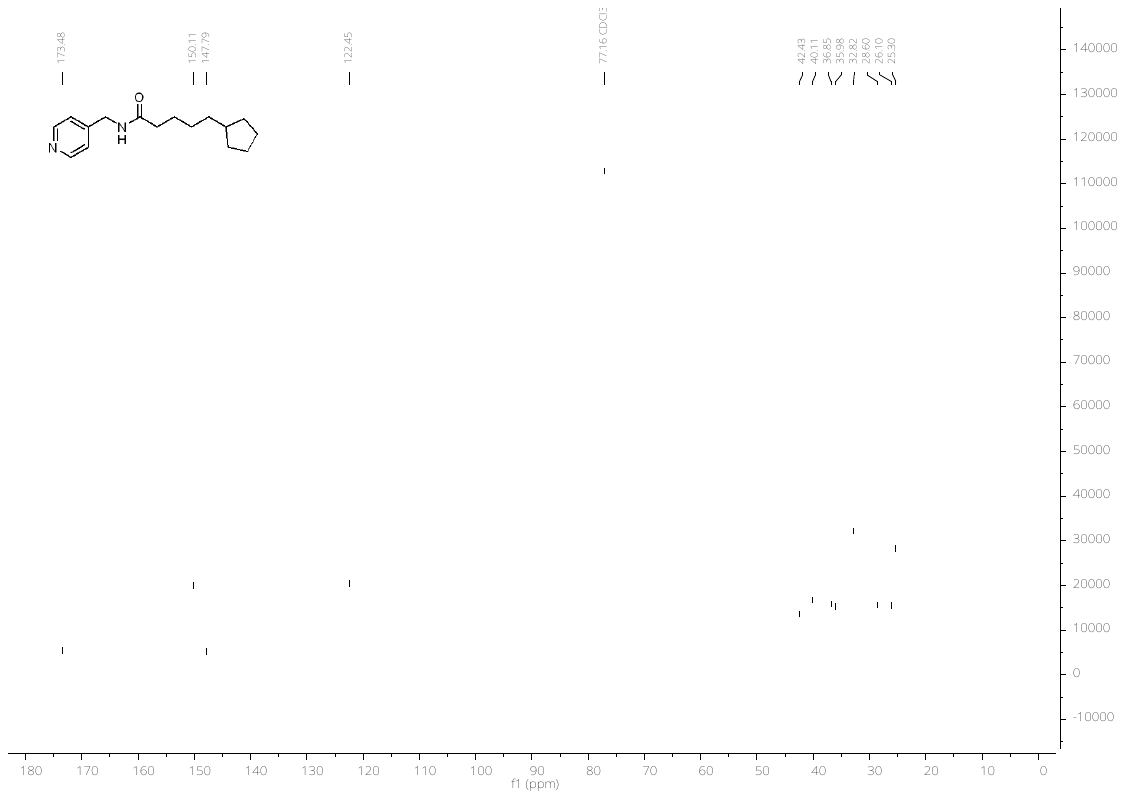

126 MHz ^13^C-NMR in CDCl_3_

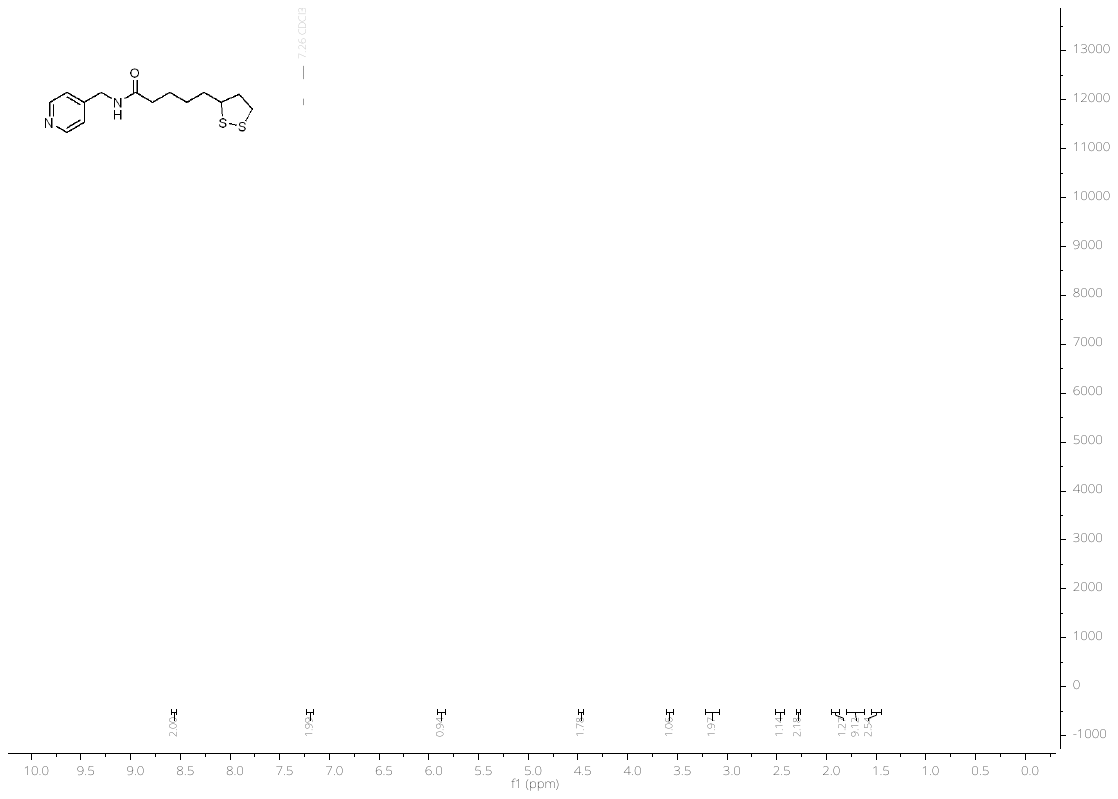

500 MHz ^1^H-NMR in CDCl_3_

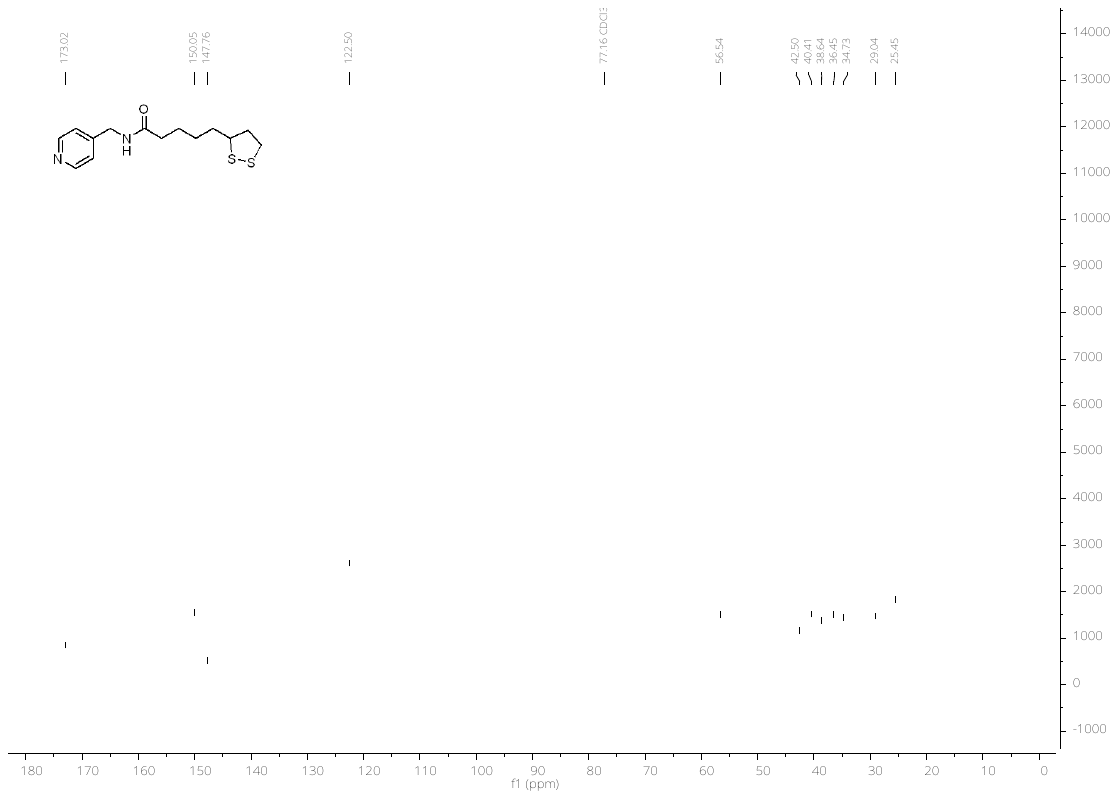

126 MHz ^13^C-NMR in CDCl_3_

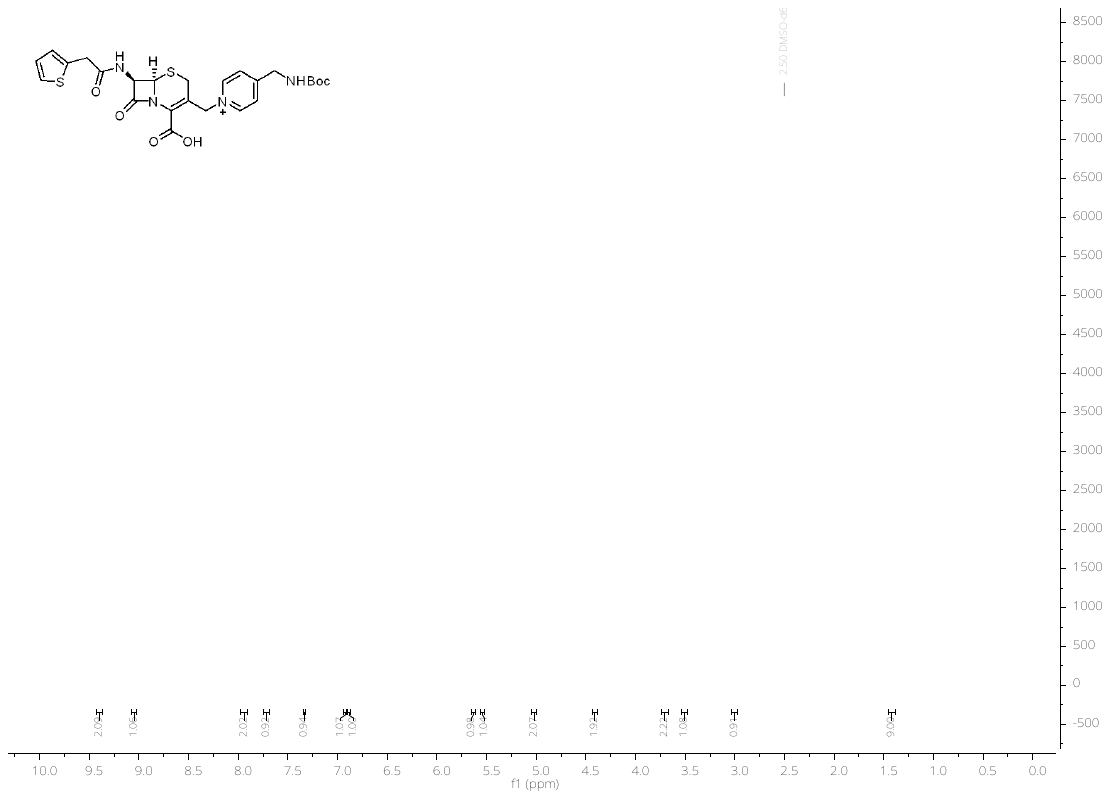

500 MHz ^1^H-NMR in DMSO-*d_6_*

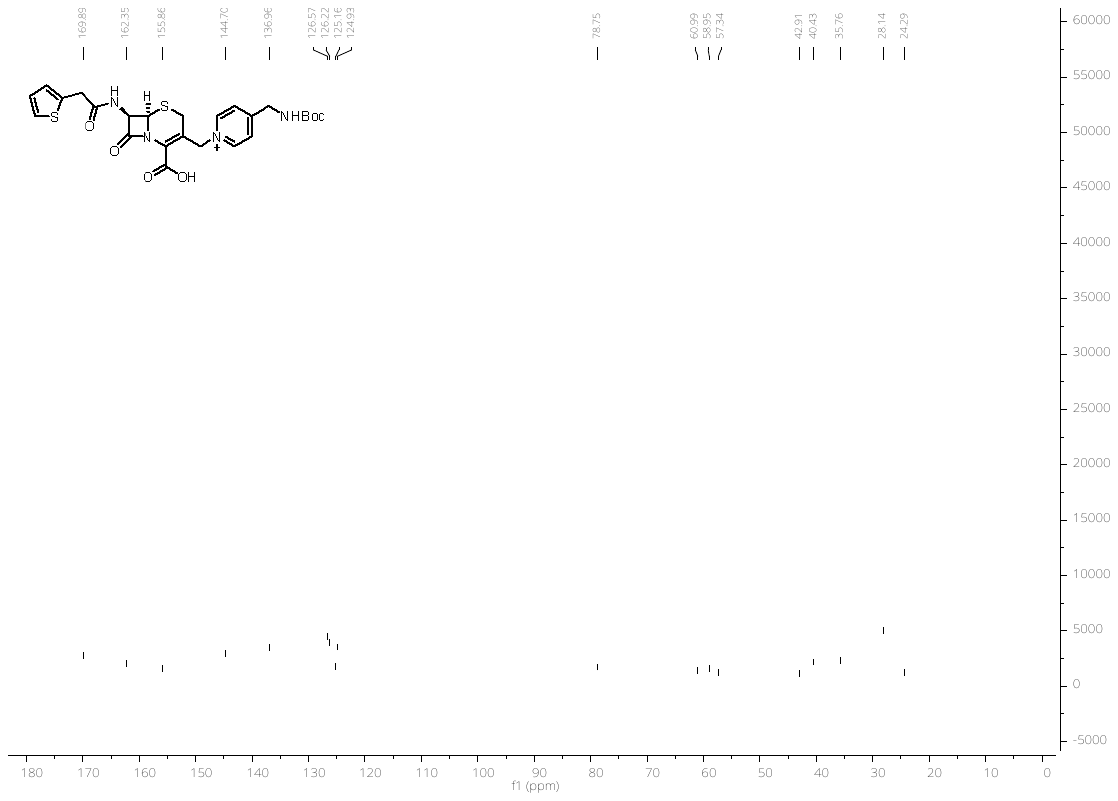

126 MHz ^13^C-NMR in DMSO-*d_6_*

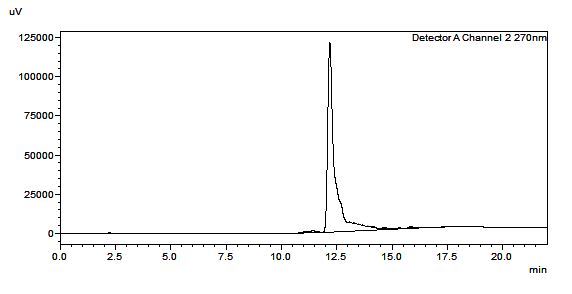

HPLC trace of cephalosporin derivative **10**

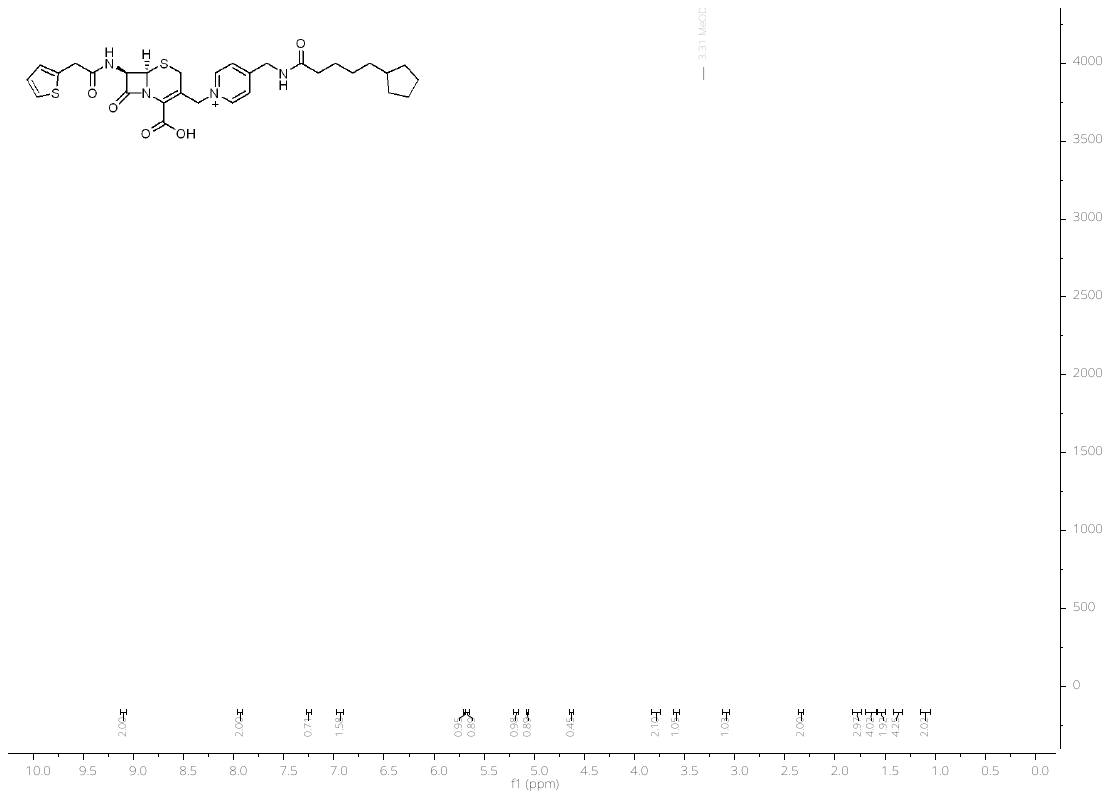

500 MHz ^1^H-NMR in MeOD of **12a**

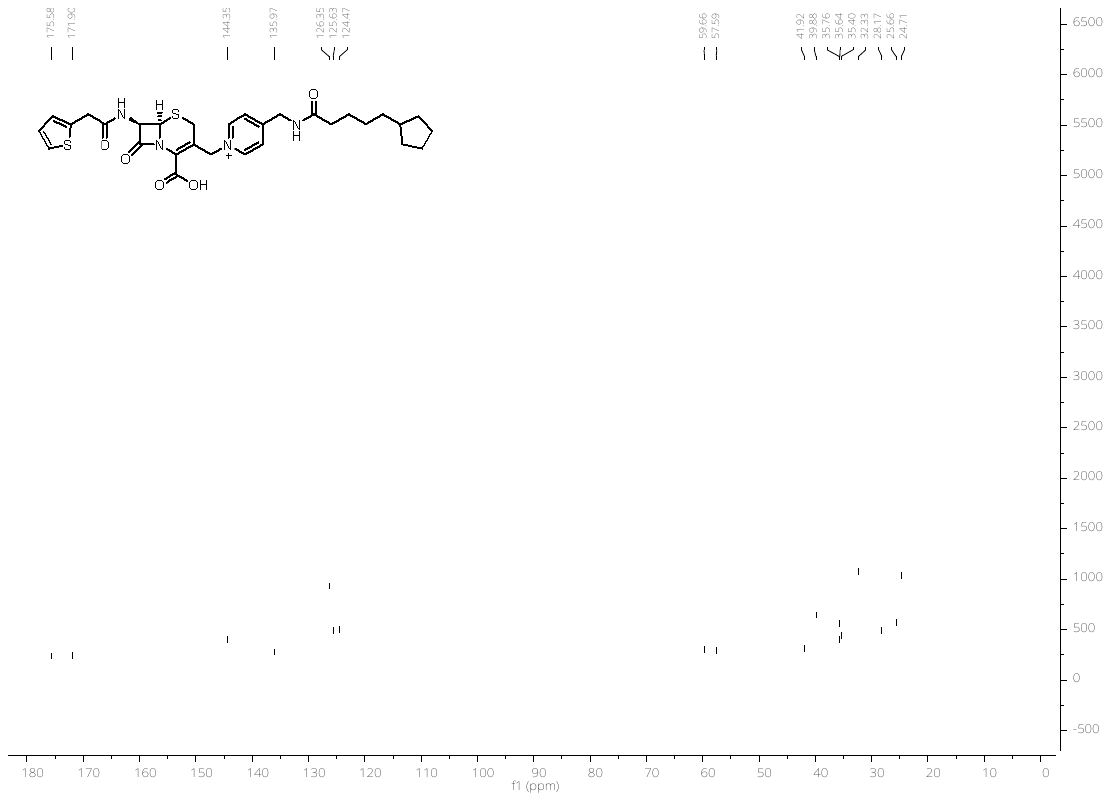

126 MHz ^13^C-NMR in MeOD of **12a**

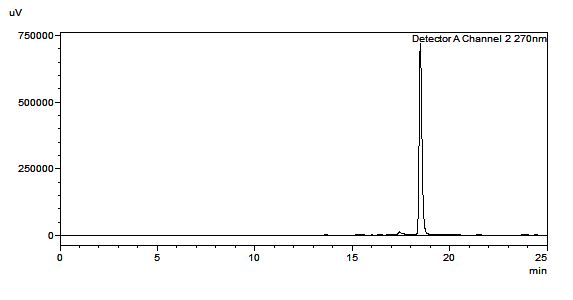

HPLC trace of compound **12a**

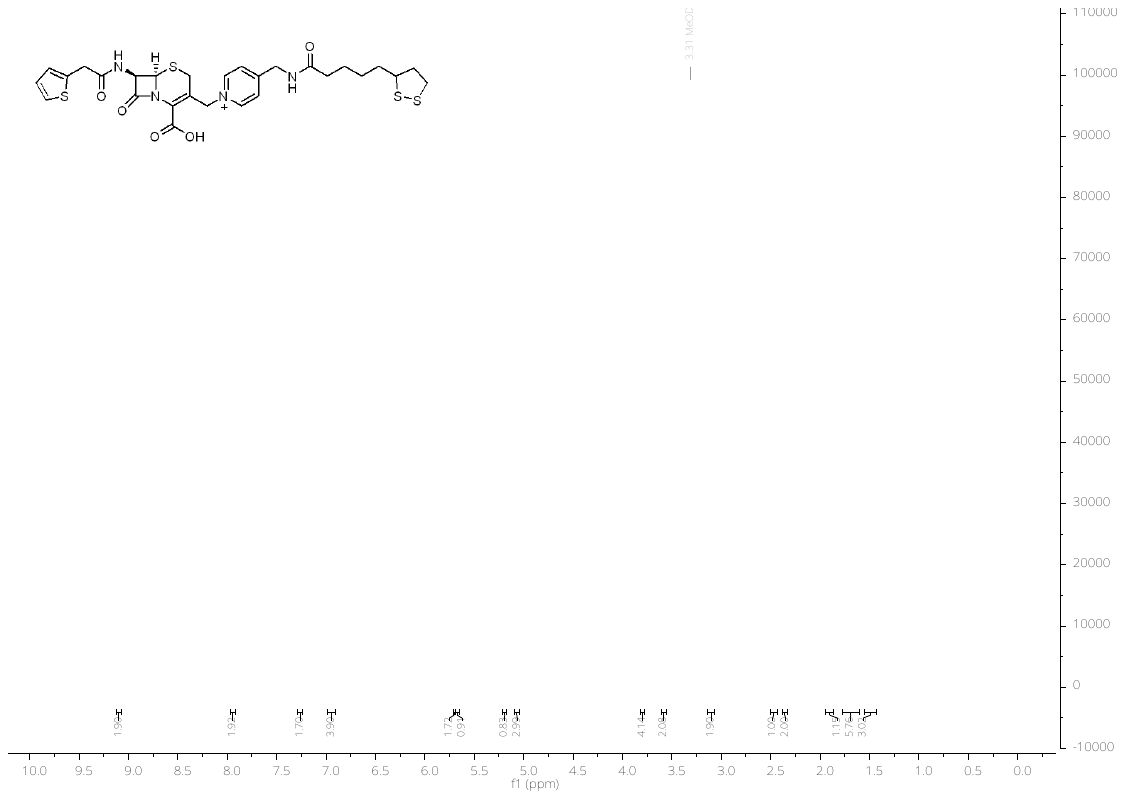

500 MHz ^1^H-NMR in MeOD of **12b**

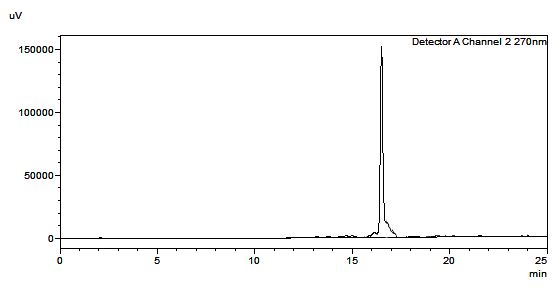

HPLC trace of compound **12b**

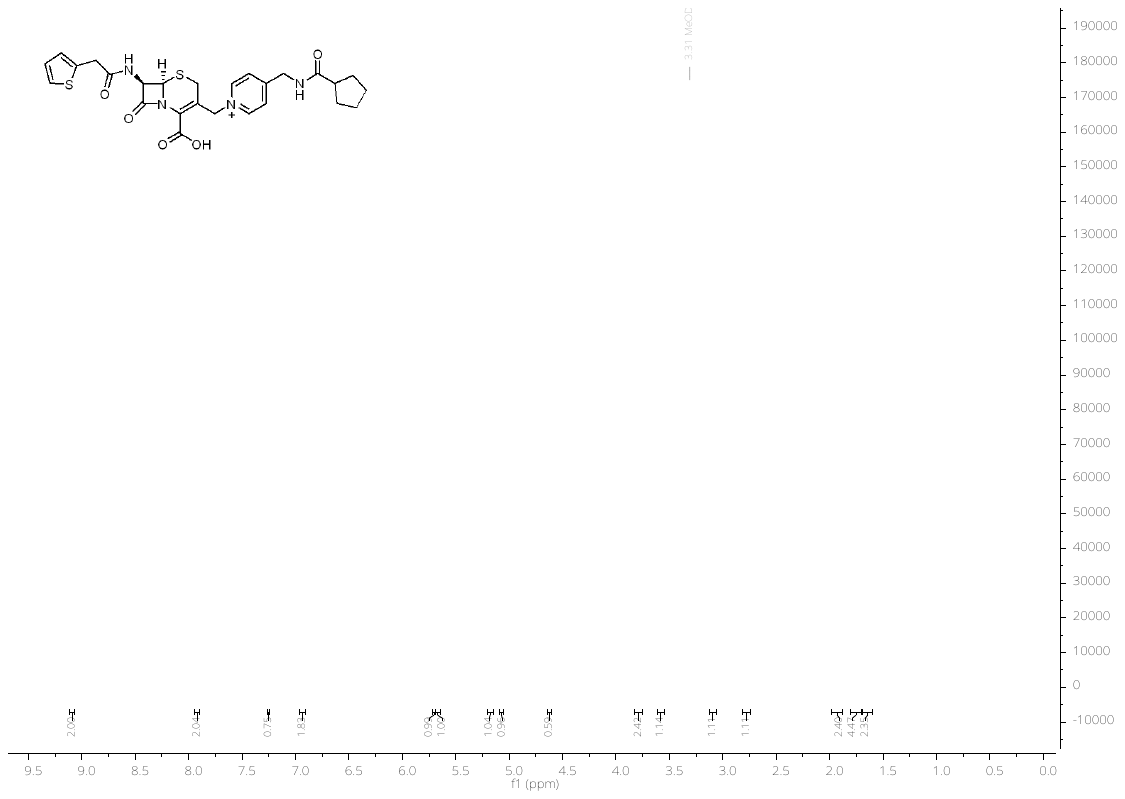

500 MHz ^1^H-NMR in MeOD of **13a**

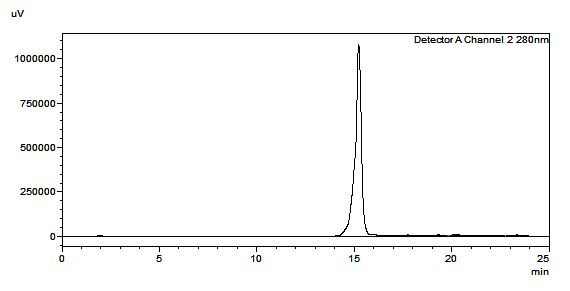

HPLC trace of the compound **13a**

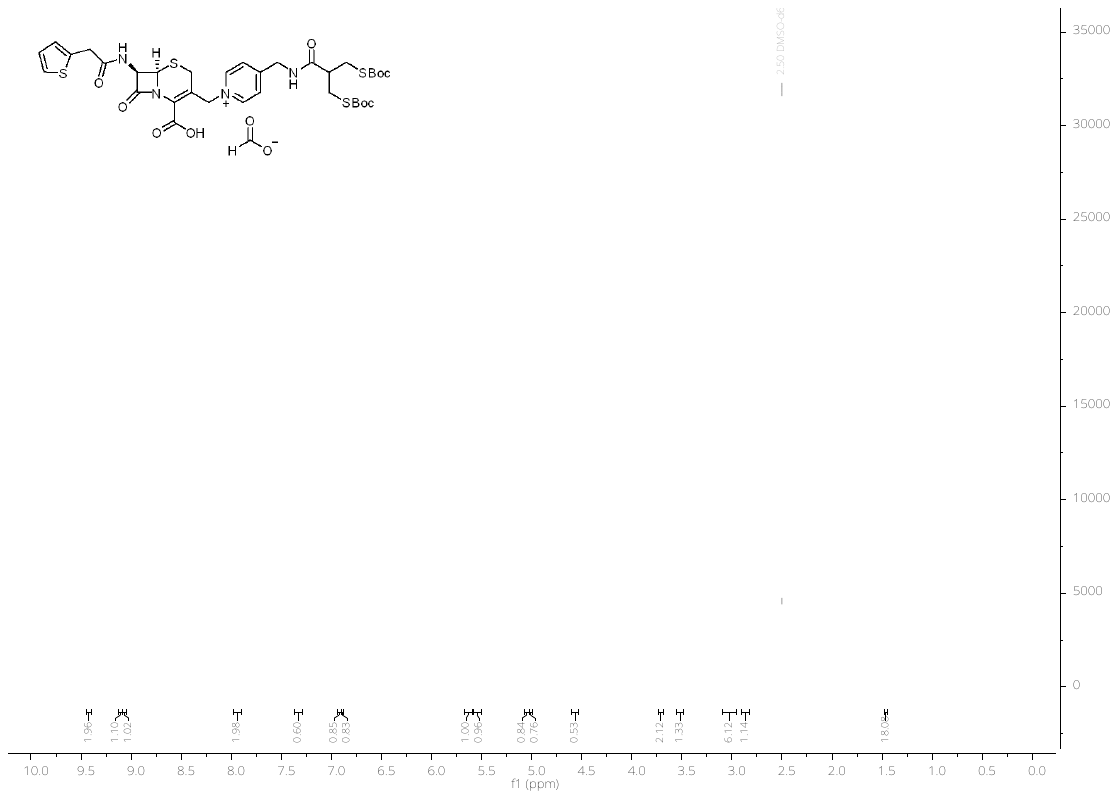

500 MHz ^1^H-NMR in DMSO-*d_6_* of **14**

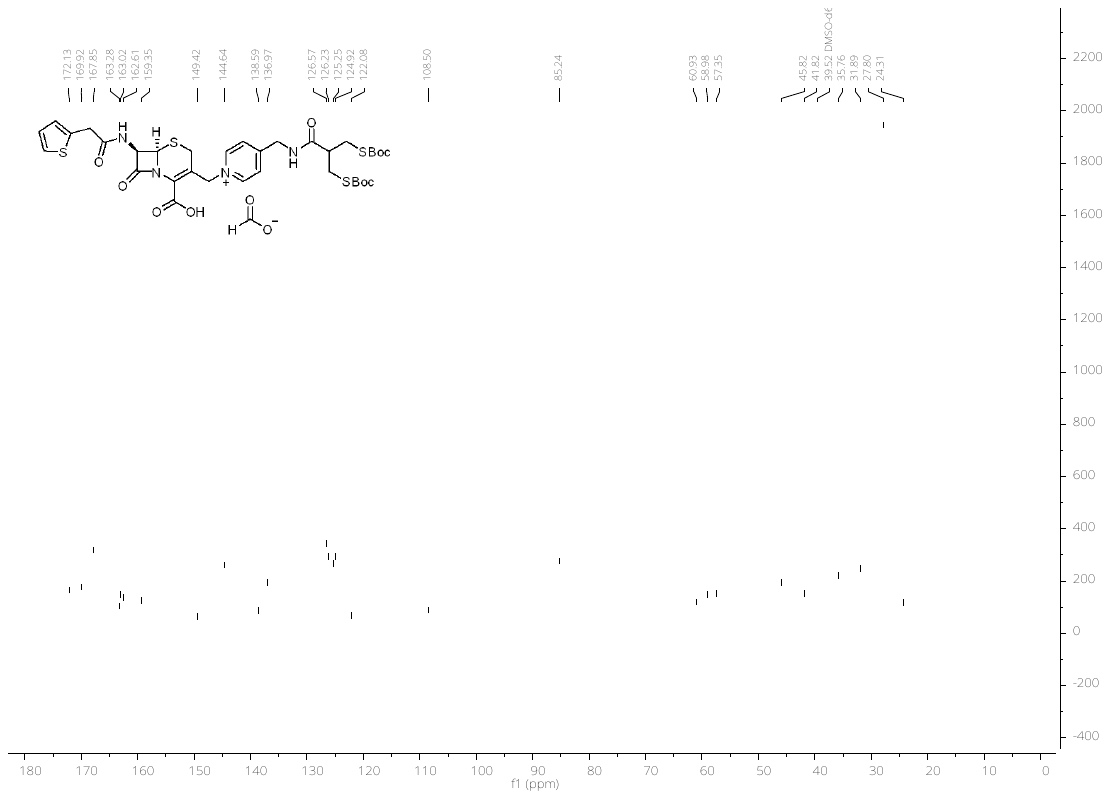

126 MHz ^13^C-NMR in DMSO-*d_6_* of **14**

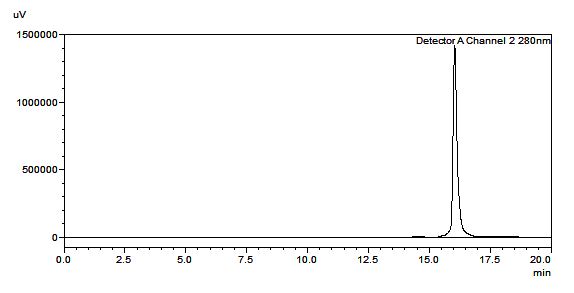

HPLC trace of the compound **14**

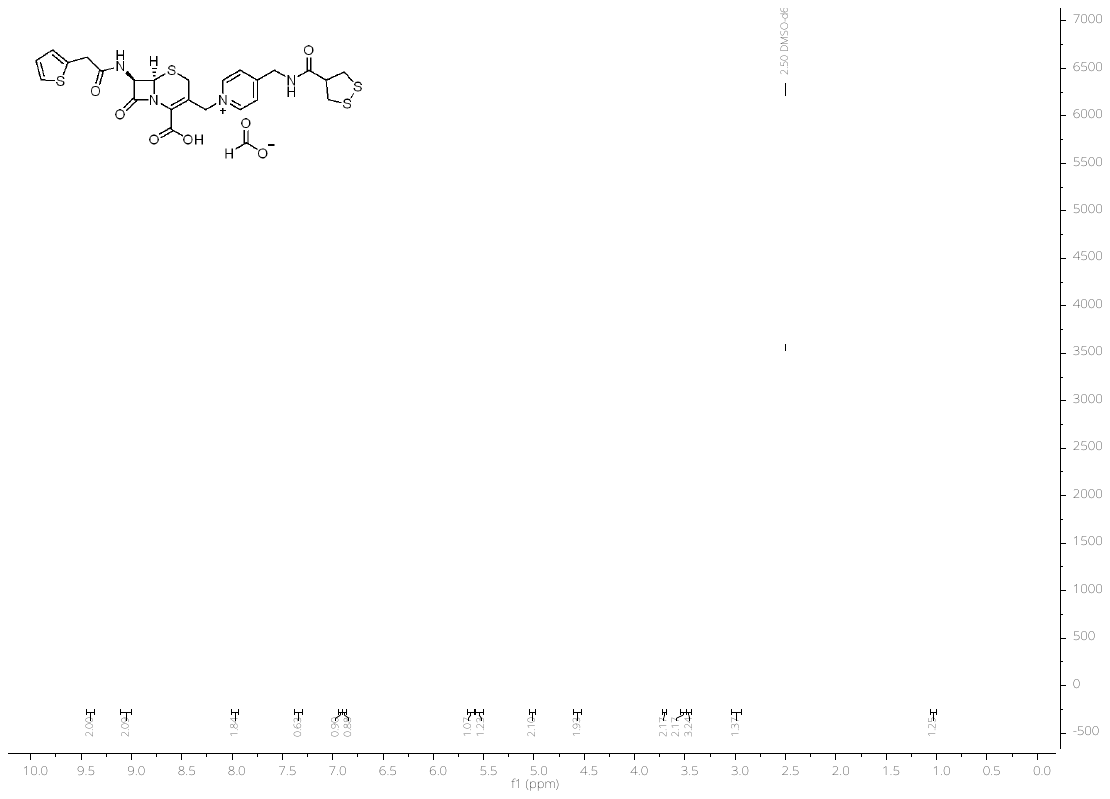

500 MHz ^1^H-NMR in DMSO-*d_6_* of **13b**

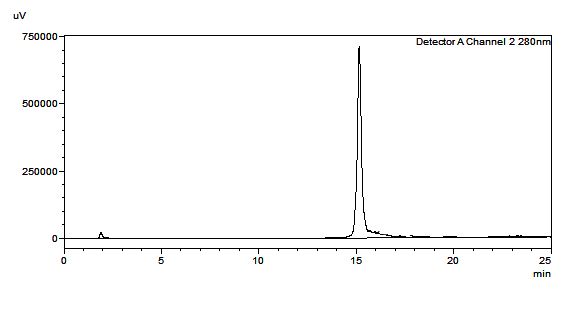

HPLC trace of the compound **13b**

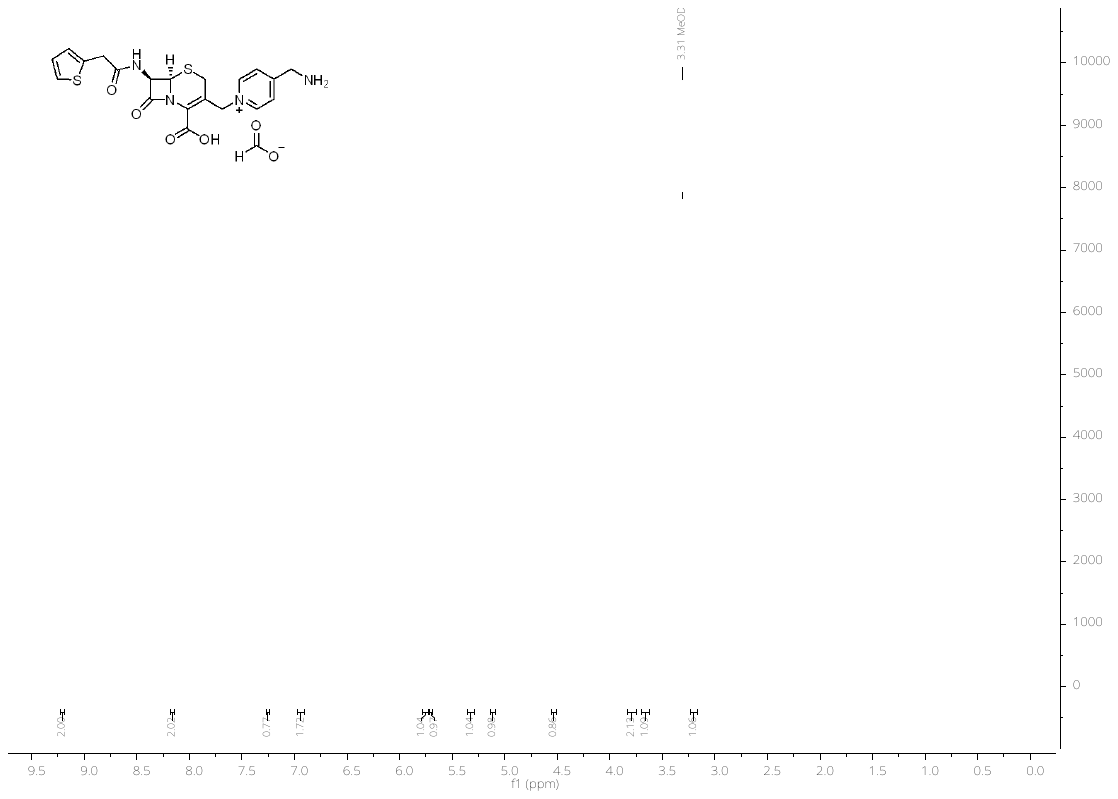

500 MHz ^1^H-NMR in MeOD of **11**

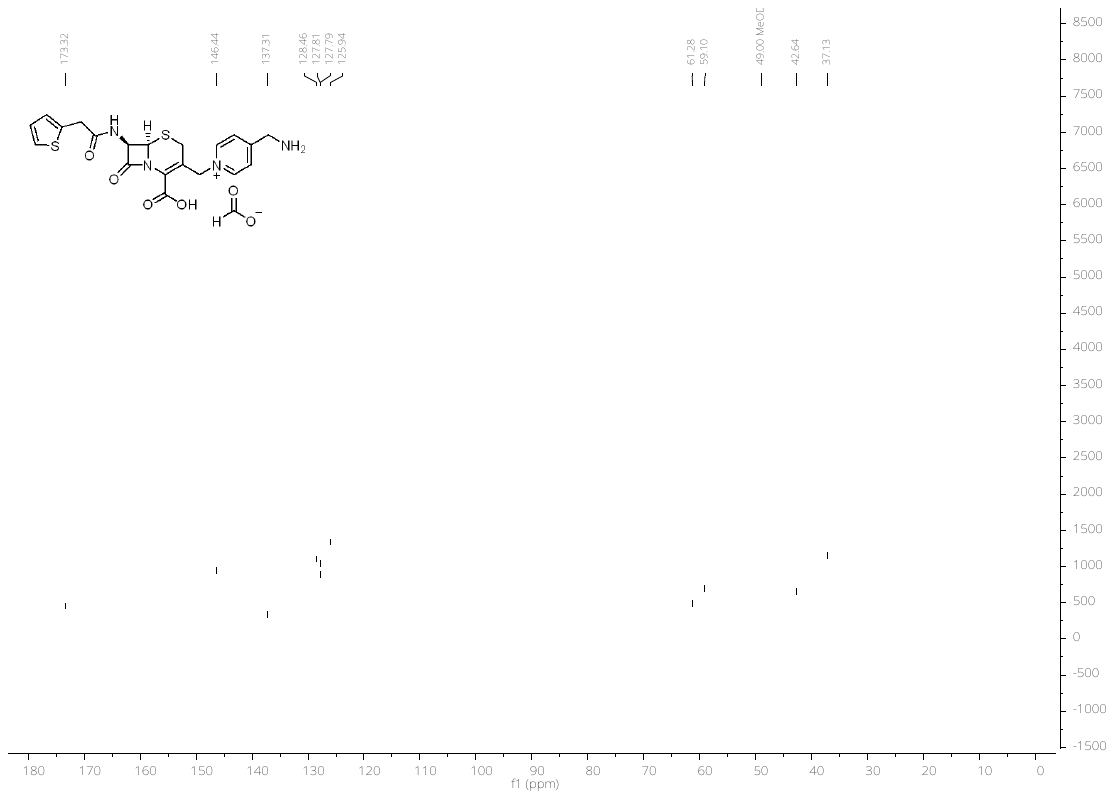

126 MHz ^13^C-NMR in MeOD of **11**

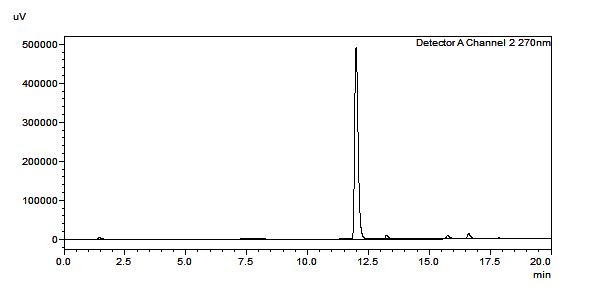

HPLC trace of the compound **11**

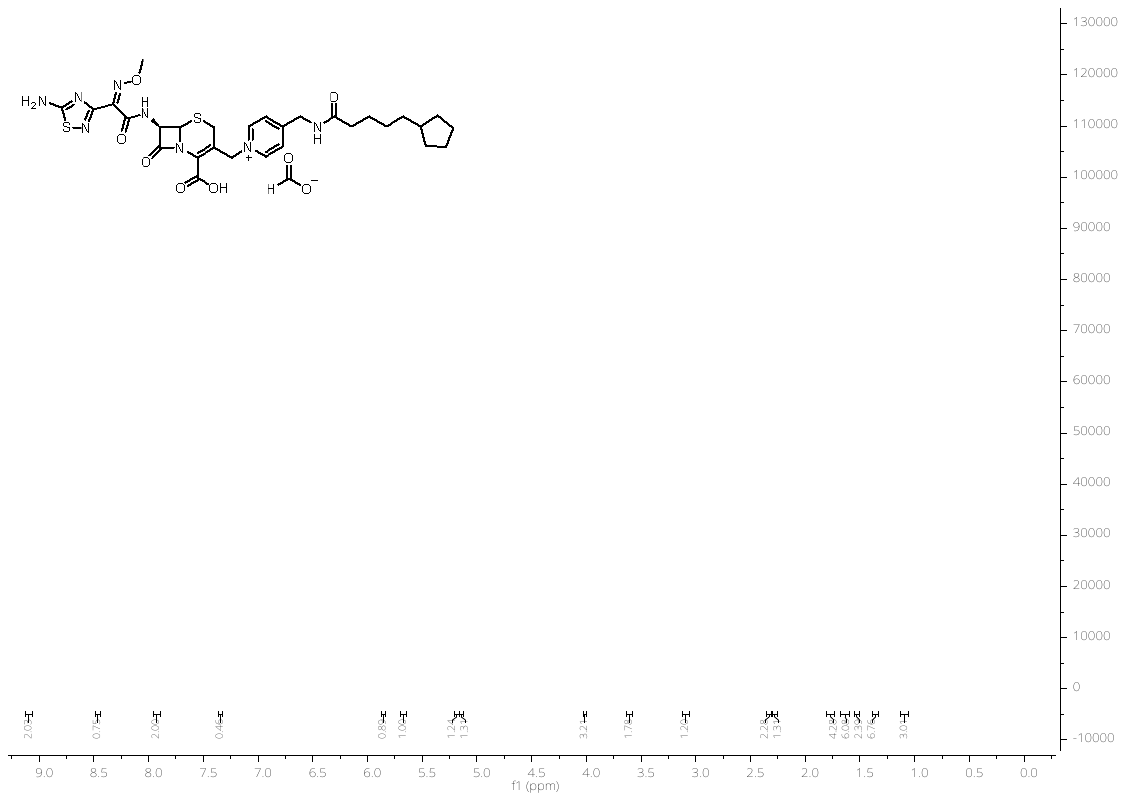

500 MHz ^1^H-NMR in MeOD of **17a**

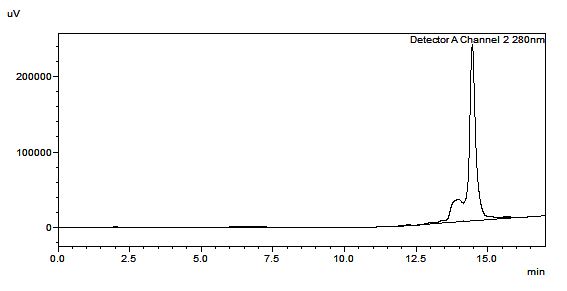

HPLC trace of compound **17a**

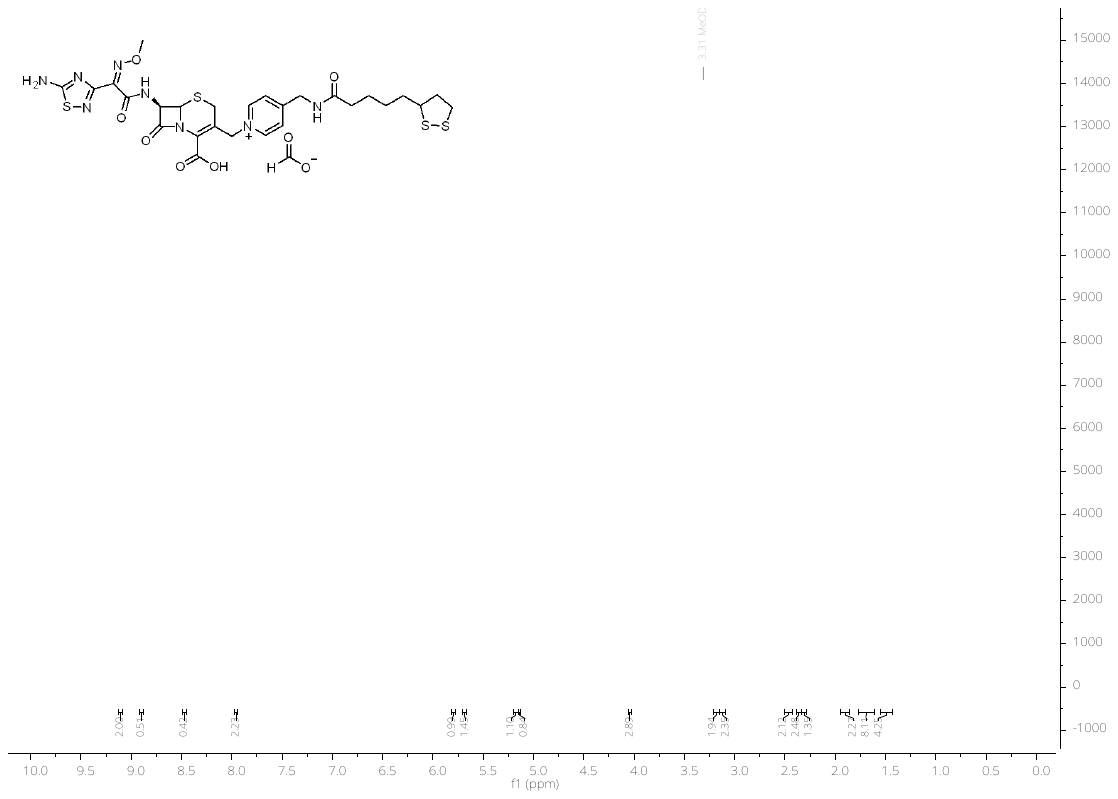

500 MHz ^1^H-NMR in MeOD of **17b**

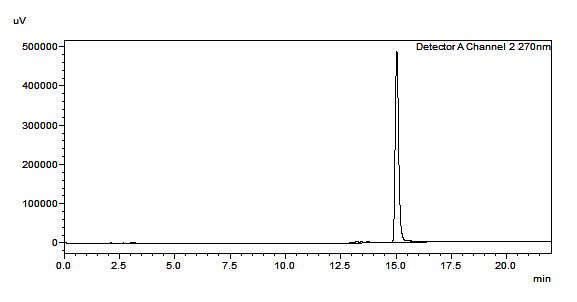

HPLC trace for compound **17b**

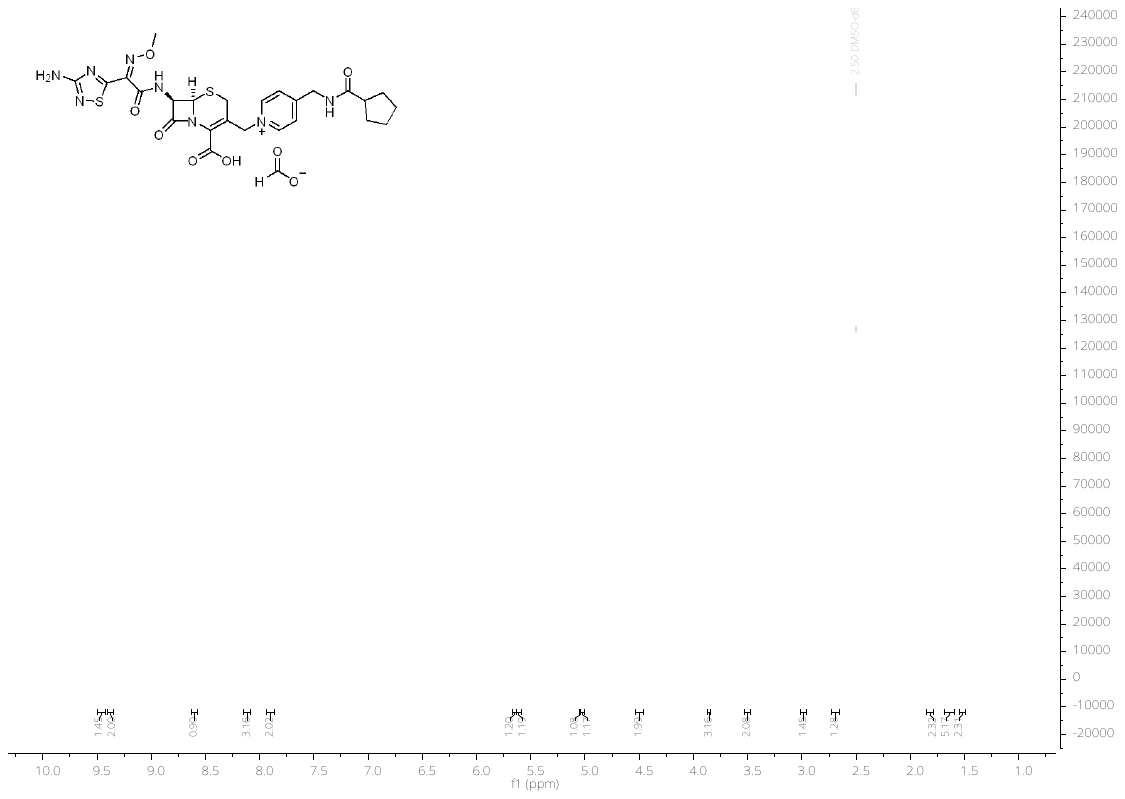

500 MHz ^1^H-NMR in DMSO-*d_6_* + 0.1%TMS of **18a**

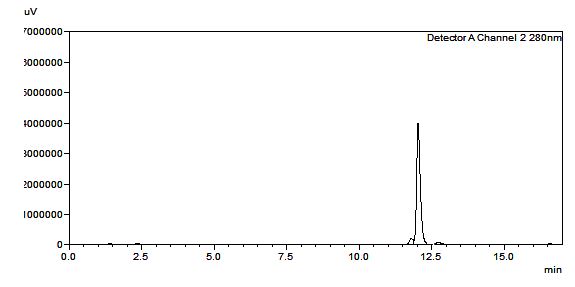

HPLC trace of compound **18a**

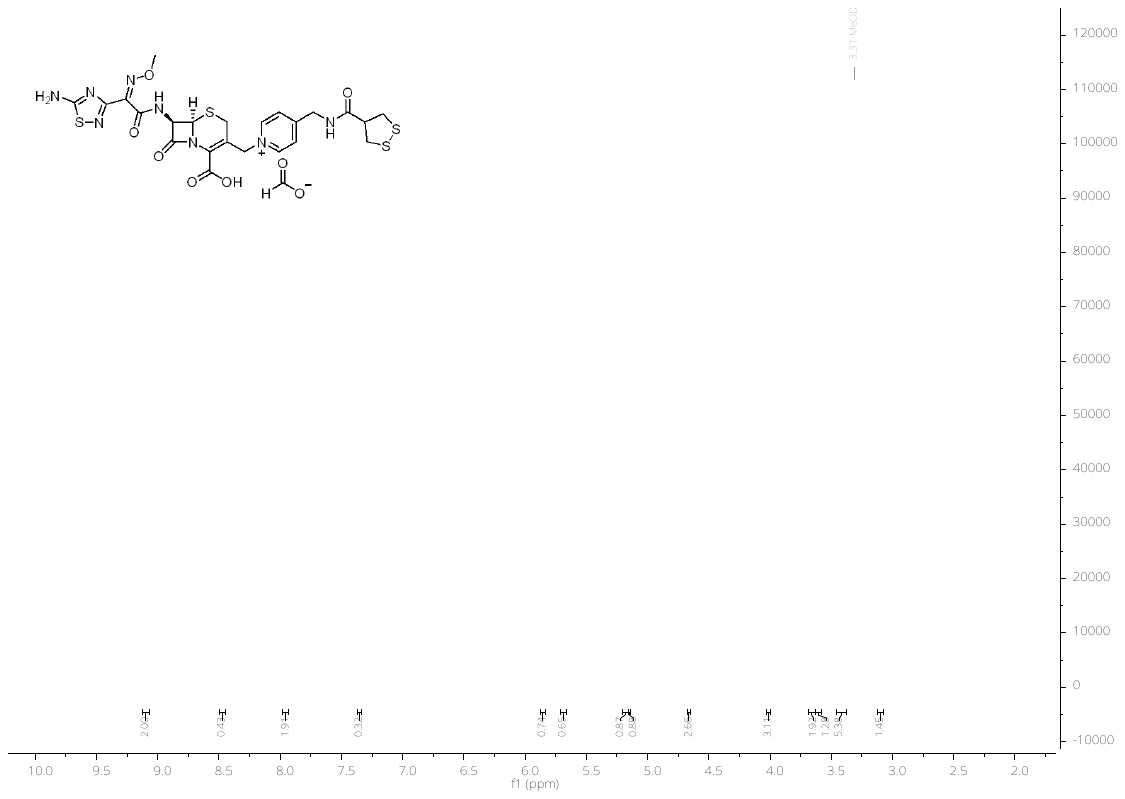

500 MHz ^1^H-NMR in MeOD of **18b**

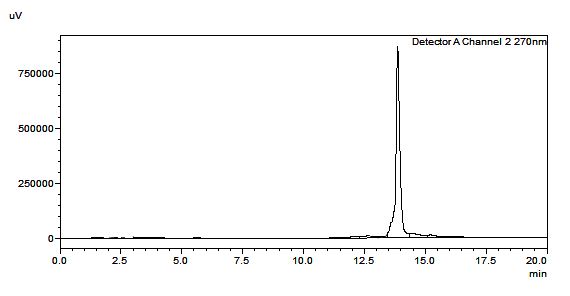

HPLC trace of compound **18b**

### III. **Microbiological assay****s**

**Minimum inhibitory concentration**

The turbidity of the overnight culture of *B. subtilis* ATCC 6633, *S. aureus* (ATCC 25923, 29213, 43300), *E. coli* (ATCC 25922, K12 MG1622), *P. aeruginosa* ATCC 27853 and PAO1 was adjusted to McFarland standard 0.5 (OD = 0.08 – 1.1), and then diluted 1 to 200 for *B. subtilis*, 1:150 for *E. coli*, and 1:100 for the *S. aureus* and *P. aeruginosa* in cation-adjusted Mueller-Hinton-II broth (MHB). The turbidity of the overnight culture *E. faecalis* ATCC 51299, was adjusted to McFarland standard 0.5 and then diluted 1 to 100 in Brain Heart Infusion Broth. The compounds were dissolved either in water or water/5% DMSO at a concentration of 1 mg/ml. In 96-wells microtiter plates, two-fold serial dilutions of the compounds (ranging from 64 μg/ml to 0.06 μg/ml) were prepared in MHB or BHI depending on strain to a final volume of 50 μl. 0.002% Tween 80 was added for testing vancomycin derivatives. The bacterial suspension (50 μl) was added into each well on the microtiter plate for inoculation, corresponding to approximately 5x10^5^ CFU/ml. The microplates were shaken (200 rcf/min) overnight at 37 °C. The MIC was determined by visual inspection of bacterial growth.

**Table S1.** MIC results for cephalosporin derivatives with thiophene side chain.

|  | MIC (µg/mL) | | | | | | |
| --- | --- | --- | --- | --- | --- | --- | --- |
|  | **3** | **11** | **10** | **13a** | **13b** | **12a** | **12b** |
| **Gram-positive strains** |  | | | | | | |
| *S. aureus* ATCC 25923 (VSSA) | 0.125 | 0.5 | 0.5-1 | 0.25 | 0.25 | 0.06 | 0.06 |
| *S. aureus* ATCC 29213 (VSSA) | 0.125 | 0.125 | 0.125 | 0.25-0.5 | 0.25 | 0.06 | 0.06 |
| *S. aureus* ATCC 43300 (MRSA) | 0.25 | 2-4 | 0.25 | 4 | 1-2 | 1-2 | 1 |
| *E faecalis* ATCC 51299 (VanB) | 16 | 32 | 32 | 32 | 32 | 8 | 8 |
| *B. subtilis* ATCC 6633 | 0.125 | 1 | 0.125 | 0.25 | 0.125 | 0.125 | 0.125 |
| **Gram-negative strains** |  | | | | | | |
| *E. coli* ATCC 25922 | 8 | 8 | 32 | 32 | 16 | >64 | >64 |
| *E. coli* K12 MG1622 | 8 | 4-8 | 32 | 32 | 8 | >64 | >64 |
| *P. aeruginosa ATCC 27853* | >64 | >64 | >64 | >64 | >64 | >64 | >64 |
| PAO1 | >64 | >64 | >64 | >64 | >64 | >64 | >64 |

**Table S2.** MIC results for cephalosporin derivatives with amino thiadiazol side chain

|  | MIC (µg/mL) | | | | |
| --- | --- | --- | --- | --- | --- |
|  | **16** | **18a** | **18b** | **17a** | **17b** |
| **Gram-positive strains** |  |  |  |  |  |
| *S. aureus* ATCC 25923 (VSSA) | 4 | 32-64 | 8-16 | 16 | 8-16 |
| *S. aureus* ATCC 29213 (VSSA) | 8 | 32-64 | 8-16 | 16 | 8-16 |
| *S. aureus* ATCC 43300 (MRSA) | 32 | >64 | 32 | 16 | 16-32 |
| *E faecalis* ATCC 51299 (VanB) | >32 | 32 | 32 | 32 | 32 |
| *B. subtilis* ATCC 6633 | 2-4 | 8-16 | 4 | 32 | 4 |
| **Gram-negative strains** |  |  |  |  |  |
| *E. coli* ATCC 25922 | 1-2 | 16-32 | 2-4 | 32-64 | 16 |
| *E. coli* K12 MG1622 | 0.25-0.5 | 8 | 1 | 32 | 64 |
| *P. aeruginosa ATCC* 27853 | 2-4 | 32-64 | 16-32 | >64 | 64 |
| PAO1 | 2-4 | 32 | 8 | >64 | 64 |

**Assessment of metabolic activity of biofilm cells by fluorescein diacetate (FDA).**

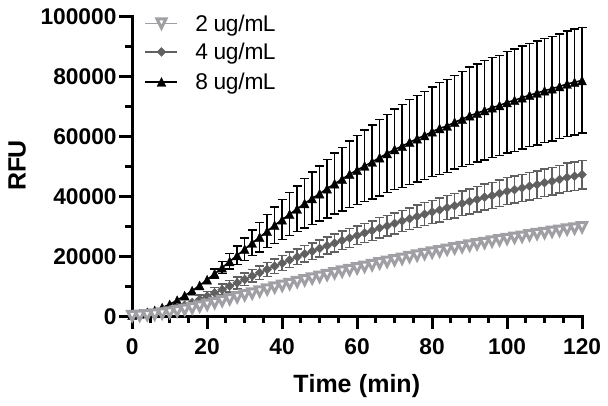

**Figure S1**. Assessment of metabolic activity of biofilm cells by FDA using three concentrations of solutions (2, 4, and 8 µg/mL): *P. aeruginosa* ATCC 27853, incubated at 37 ^o^C. Relative fluorescence measurements were taken every 2 min for 2 h. Error bars represent standard deviation of two independent experiments in triplicate.

**Calculation of data viability for the experiments of metabolic activity of biofilm cells**

Statistical quality parameters were calculated according to the method described before.^1,2^

The data was combined and the Mean value and SD was calculated by Prism from 6 data points.

Then obtained data was used for determining assay performance or sensitivity

X*_max_* – means of maximal signal

X*_min_* – means of minimal signal

SD*_max_* – the standard deviation of maximal signal

SD*_min_* – the standard deviation of minimal signal

Signal to noise $S:N= \frac{X_{max}- X_{min}}{\sqrt{{{SD}_{max}}^{2}- {{SD}_{min}}^{2}}}$

Signal to background $S:B= \frac{X_{max}}{X_{min}}$

Signal window $SW= \frac{X_{max}-X_{min}-6({SD}_{max}-{SD}_{min})}{{SD}_{max}}$

Z’ factor $Z^{'}=1-\frac{6{SD}_{max}-6{SD}_{min}}{\left| X_{max}-X_{min} \right|}$

**Table S3.** Calculation of quality parameters for metabolic activity of biofilm cells assay

| Bacterial  strains | FDA  concentration  (µg/mL) | Time (min) | 10 | 20 | 30 | 40 | 50 | 60 | 70 | 80 | 90 | 100 | 110 | 120 |
| --- | --- | --- | --- | --- | --- | --- | --- | --- | --- | --- | --- | --- | --- | --- |
| *P. aeruginosa*  ATCC 27853 | 2 | S/N | 4.03 | 8.10 | 13.30 | 14.51 | 14.68 | 15.46 | 16.07 | 17.94 | 18.01 | 18.67 | 20.27 | 20.49 |
|  |  | S/B | 29.60 | 55.42 | 66.85 | 69.32 | 67.14 | 64.38 | 60.82 | 56.80 | 52.89 | 49.87 | 46.75 | 43.71 |
|  |  | SW | 1.05 | 5.11 | 10.32 | 11.58 | 11.75 | 12.54 | 13.15 | 15.04 | 15.13 | 15.80 | 17.42 | 17.65 |
|  |  | Z' | -0.49 | 0.26 | 0.55 | 0.58 | 0.59 | 0.61 | 0.62 | 0.66 | 0.66 | 0.67 | 0.69 | 0.70 |
|  | 4 | S/N | 2.83 | 4.07 | 5.80 | 7.04 | 7.79 | 8.63 | 8.67 | 8.88 | 8.94 | 9.06 | 9.43 | 9.63 |
|  |  | S/B | 27.65 | 52.28 | 63.66 | 64.70 | 62.25 | 57.75 | 53.65 | 49.36 | 45.78 | 42.47 | 39.40 | 36.58 |
|  |  | SW | -0.12 | 1.11 | 2.85 | 4.12 | 4.88 | 5.75 | 5.80 | 6.04 | 6.11 | 6.26 | .6.66 | 6.88 |
|  |  | Z' | -1.14 | -0.48 | -0.04 | 0.14 | 0.22 | 0.29 | 0.29 | 0.30 | 0.30 | 0.31 | 0.33 | 0.34 |
|  | 8 | S/N | 13.06 | 10.87 | 5.84 | 4.80 | 4.38 | 4.16 | 4.10 | 4.05 | 4.05 | 4.11 | 4.24 | 4.36 |
|  |  | S/B | 28.84 | 61.65 | 77.90 | 82.43 | 79.17 | 73.78 | 68.17 | 61.59 | 56.31 | 50.97 | 46.32 | 42.15 |
|  |  | SW | 10.17 | 7.89 | 2.86 | 1.82 | 1.40 | 1.18 | 1.12 | 1.07 | 1.07 | 1.14 | 1.27 | 1.39 |
|  |  | Z' | 0.53 | 0.44 | -0.03 | -0.25 | -0.37 | -0.44 | -0.47 | -0.48 | -0.48 | -0.46 | -0.42 | -0.38 |

**Table S4.** MBRC results for cephalosporin derivatives

| Bacterial strain | Antibiotic | MIC (µg/mL) | MBRC (µg/mL) | |
| --- | --- | --- | --- | --- |
|  |  |  | Resazurin  (decrease in RFU) | CFU/mL  (growth inhibition) |
| *S. aureus* ATCC 29213 | **3** | 0.125 | 0.25 (100%) | 0.25 (100%) |
|  | **10** | 0.125 | 0.5 (98%) | 0.5 (100%) |
|  | **11** | 0.125 | 0.5 (99%) | 0.5 (100%) |
|  | **12a** | 0.06 | 0.25 (98%) | 0.25 (100%) |
|  | **12b** | 0.06 | 0.5 (99%) | 0.5 (100%) |
|  | **13a** | 0.25-0.5 | 0.5 (100%) | 0.5 (100%) |
|  | **13b** | 0.5 | 0.5 (99%) | 0.5 (100%) |
|  | **16** | 8 | 16 (99%) | 16 (100%) |
|  | **17a** | 16 | 16 (100%) | 16 (100%) |
|  | **17b** | 8-16 | 16 (100%) | 16 (100%) |
|  | **18a** | 32-64 | >32 | >32 |
|  | **18b** | 8-16 | 16 (100%) | 16 (100%) |
| *S. aureus* ATCC 43300  (MRSA) | **3** | 0.25 | 1 (97%) | 1 (100%) |
|  | **10** | 0.25 | 4 (99%) | 4 (100%) |
|  | **11** | 2-4 | 2 (98%) | 2 (100%) |
|  | **12a** | 1-2 | 1 (99%) | 1 (100%) |
|  | **12b** | 1 | 2 (100%) | 2 (100%) |
|  | **13a** | 4 | 4 (100%) | 4 (100%) |
|  | **13b** | 1-2 | 4 (100%) | 4 (100%) |
|  | **16** | 32 | 16 (92%) | 16 (100%) |
|  | **17a** | 16 | 16 (99%) | 16 (100%) |
|  | **17b** | 16-32 | 32 (98%) | 32 (100%) |
|  | **18a** | >64 | 64 (87%) | 64 (99%) |
|  | **18b** | 32 | 32 (100%) | 32 (100%) |
| *E. faecalis* ATCC 51299  (VanB) | **12a** | 8 | 8 (99%) | 8 (100%) |
|  | **12b** | 8 | 16 (97%) | 16 (100%) |
|  | **13a** | 32 | 16 (98%) | 16(100%) |
|  | **13b** | 32 | 16 (96%) | 16 (100%) |
| *P. aeruginosa*  ATCC 27853 | **16** | 2-4 | 8 (91%) | 8 (99%) |
|  | **18a** | 32-64 | 64 (99%) | 64 (100%) |
|  | **18b** | 16-32 | 16 (96%) | 16 (100%) |
